# Task-independent metrics of computational hardness predict human cognitive performance

**DOI:** 10.1101/2021.04.25.441300

**Authors:** Juan P. Franco, Karlo Doroc, Nitin Yadav, Peter Bossaerts, Carsten Murawski

**Affiliations:** Brain, Mind and Markets Laboratory, Department of Finance, The University of Melbourne, Parkville, Victoria 3010, Australia; Florey Institute of Neuroscience and Mental Health, Parkville, Victoria 3010, Australia

## Abstract

The survival of human organisms depends on our ability to solve complex tasks in the face of limited cognitive resources. However, little is known about the factors that drive the complexity of those tasks. Here, building on insights from computational complexity theory, we quantify the computational hardness of cognitive tasks using a set of task-independent metrics related to the computational resource requirements of individual instances of a task. We then examine the relation between those metrics and human behavior and find that they predict both time spent on a task as well as accuracy in three canonical cognitive tasks. Our findings demonstrate that performance in cognitive tasks can be predicted based on generic metrics of their inherent computational hardness.

**Teaser:** The ability of humans to solve cognitive tasks is affected by generic mathematical properties of problems related to their computational complexity.

## Introduction

People are remarkably flexible in their ability to adapt to changes in their environment and their ability to solve a vast range of complex cognitive tasks. This is even though we are limited in our computational resources such as time and memory (*1*–*3*). Computational limitations are particularly relevant since many of the daily tasks people face involve solving computational problems that are considered computationally hard (*4*). Everyday life examples of deceptively simple yet actually hard problems include attention gating, task scheduling, shopping, routing, bin packing, and game play (*4*, *5*). This raises the question of how hardness of problems interacts with human computational limitations and how it affects the ability of humans to solve day-to-day cognitive tasks.

Here, we draw on computational complexity theory to examine human *performance* on a set of canonical cognitive tasks and test whether it is related to a set of task-independent measures of computational complexity. Specifically, we aim to show that generic properties of individual instances of a task, associated with computational hardness, predict human behavior. Our goal is to characterize the computational hardness of problems in a way that is independent of the particular problem. This would mark a step towards a general theory linking properties of the task environment to human performance. Such theory could provide policy prescriptions where the cognitive demands of a task substantially exceed human capacities without needing to take a stance on the strategies people employ. As such, this research program could inform public policy over the computational hardness of products such as insurance and financial products. Moreover, it could inform the development of artificial intelligence systems adjusted to inform human decision-makers when computational requirements exceeds capacities.

To date, research on the effect of computational hardness of instances of problems on human performance lacks generality. Most studies are based on a problem- or algorithm-specific approach, that is, they are based on measures of hardness specific to a particular problem (*6*–*9*) or a particular solving strategy (*10*–*12*). Thus, those findings may not generalize to other problems or across strategies. Recent theoretical advances in computer science and statistical physics provide a framework, referred to as typical-case complexity (TCC), that addresses this issue.

TCC is concerned with the average computational hardness of random instances of a computational problem, linking structural properties of those instances to their computational complexity, independent of a particular computational model (*13*–*18*). This work has identified computational ‘phase transitions’, which resemble phase transitions in statistical physics and which are related to computational hardness of instances. Such phase transitions have been found in a number of canonical NP-complete problems (i.e., problems that are both in NP and NP-hard) (*13*–*18*), including the knapsack problem (*18*), the traveling salesperson problem (*13*) and the K-SAT problems (Boolean satisfiability problems) (*14*, *19*, *20*), among others. These are problems where a candidate solution can be verified in *polynomial time* (solve-time of an algorithm grows at most polynomially in the size of the problem), but no algorithms are currently known that can reliably compute the solution in polynomial time. This program has led to a deeper understanding of computational hardness by relating it to structural properties of instances, as well topological properties of the solution space. Importantly, it has identified that the hardness of an instance is related to a generic and intrinsic property of an instance: constrainedness (see Fig 1). Specifically, TCC captures the average computational hardness of random instances for a particular level of constrainedness.

**Figure 1:**
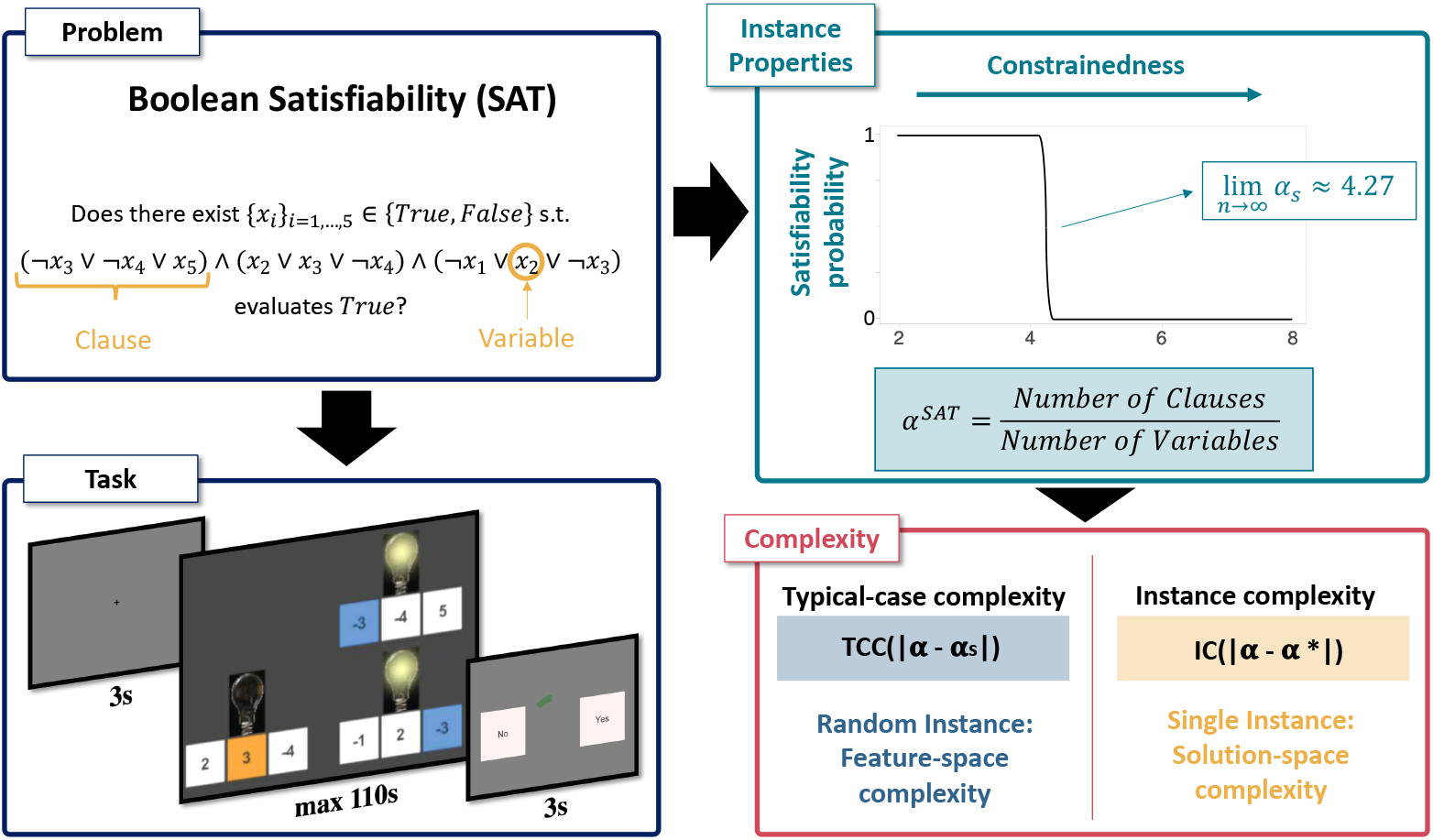
3SAT problem, complexity metrics and experimental design. The problem. The aim is to determine whether a Boolean formula is *satisfiable*. **The task.** The Boolean formula is represented with a set of light bulbs (clauses), each of which has three switches underneath (literals) that are characterized by a positive or negative number. The number on each switch represents the variable number, which can be turned on or off (TRUE or FALSE). The aim is to determine whether there exists a way of turning on and off variables such that all the light bulbs are turned on (formula evaluates TRUE). **Instance properties.** The constrainedness of the problem (*α*) is captured by the ratio of clauses to variables. This parameter characterizes the probability that a random instance of the problem is satisfiable. In the limit this probability undergoes a phase transition around the satisfiability threshold (*α*_*s*_). **Complexity metrics.** Instances near this threshold are on average harder to solve than instances further away. Average hardness is captured by typical-case complexity (TCC). This metric can be estimated entirely from the features of the problem (feature-space) without the need to solve the problem. Alternatively, instance complexity (IC) can be estimated from features of the solution-space. IC is defined as the difference between the constrainedness of the instance (*α*) and *α*\*, the maximum number of clauses that can be satisfied divided by the number of variables in the instance.

This program has aimed to characterize the computational hardness of an instance without the need to solve the problem. As such, it has been particularly useful in predicting patterns on the performance of algorithms on random instances (*20*, *21*). However, this approach, by construction, only captures the average expected computational hardness for a collection of random instances with a fixed level of constrainedness. It does not characterize complexity at the level of an individual instance. Here, we apply an extension of this framework to study computational hardness of individual instances of problems employing a related metric: instance complexity (IC; Fig 1).

A recent study applied this framework to study human behavior in the knapsack problem, a ubiquitous combinatorial optimization problem (*22*). The study found that both time-on-task and the ability to solve an instance were related to structural properties of instances, and in particular to TCC and IC. An important question that arises is whether these findings on human performance generalize to other problems. If they do, then these properties related to computational complexity would be prime candidates for generic measures of computational hardness of human cognition, in a similar way that statistics like mean, variance and kurtosis serve as generic measures of randomness of a task (*23*, *24*).

Using a behavioral experiment, we study the relation between a set of problem-independent measures of instance complexity and human performance in two canonical NP-complete computational problems. Overall, these types of problems underlie many day-to-day cognitive tasks such as shopping, bin packing, way-finding and task scheduling (*4*, *5*). We specifically tested human performance in the Boolean satisfiability problem (3SAT) and the traveling salesperson problem (TSP), and compared these results to those previously obtained for the 0-1 knapsack problem (KP) (*22*), to test their generalizability across NP-complete problems. While the SAT task is an abstract logical problem, TSP is presented as a visual navigation task and the KP is associated with arithmetic operations. We show that the metrics of complexity affect performance similarly across the three problems, despite them being diverse in nature.

## Results

Each participant solved one of three problems: either 72 instances of the TSP task, 64 instances of the 3SAT or 72 instances of the KP. TSP is the problem of determining whether there exists a path of a particular length (or shorter), connecting a set of cities (Fig 5b). 3SAT is the problem of determining whether a set of variable configurations (true/false) exist that render a set of clauses true (Fig 1). And KP is the problem of determining whether there exists a subset of items with differing values and weights exceeding a minimum total value while not exceeding a maximum total weight (Fig 5c). All three problems are *decision problems*, that is, problems whose answer is either ‘yes’ or ‘no’. If there exists a configuration of variables such that the solution of the instance is ‘yes’, the instance is called *satisfiable* and *unsatisfiable* otherwise.

Instances varied in their computational hardness (see Materials and Methods). Both TSP and 3SAT were self-paced (with maximum time limits per trial), while the KP was not. Results for KP have previously been reported elsewhere and are included here for comparison only (*22*).

### Summary statistics

We used two metrics of performance: accuracy and time-on-task. Accuracy was measured as a binary outcome, depending on whether a participant’s response was correct or not. Time-on-task was measured as the amount of seconds a participant spent on the problem screen before advancing to the response screen. These two measures, accuracy and time-on-task, capture two different dimensions of an agent’s problem-solving process. Time-on-task is a measure of the resources deployed to solving the task while accuracy is a measure of efficacy of effort. It is worth noting that the time-on-task analysis was not performed for the KP, since this task was not self-paced.

In the TSP, all instances had 20 cities and a time limit of 40 s. The number of cities and time limit were selected, based on pilot data, to ensure that the task was neither too difficult nor too easy (see Materials and Methods). Mean *human accuracy*, measured as the proportion of trials in which a correct response was made, was 0.85 (min = 0.76, max = 0.93, *SD* = 0.05). Participants’ average time spent on an instance was 32.2 s and ranged from 19.9 s to 39.2 s (SD = 5.2). Accuracy did not vary during the course of the task, but time-on-task decreased as the task progressed (S3 Appendix).

All instances of the 3SAT task had 5 variables and a time limit of 110 s. Similar to TSP, the number of variables and time limit were selected, based on pilot data, to target a specific average accuracy (≈ 85%; see Materials and Methods). Mean *human accuracy* was 0.87 (min = 0.75, max = 0.98, *SD* = 0.06). The average time spent on an instance varied from a minimum of 15.9 s to a maximum of 104.3 s (mean = 60.2, SD = 18.7). Similar to TSP, accuracy did not vary during the course of the task, but participants tended to spend less time on a trial as the task progressed (S3 Appendix). Henceforth, for the statistical analyses, time-on-task is measured as a proportion of the maximum time allotted on each trial.

In the KP decision task implemented by Franco et al. (*22*), all instances had 6 items. This task was not self-paced, that is, participants had exactly 25 seconds to solve each instance and could not skip to the response screen before the time ended. Mean *human accuracy* was 83.1% (min = 0.56, max = 0.9, *SD* = 0.08). Like in the other two tasks, accuracy did not vary during the course of the task (S3 Appendix).

### Feature-space complexity metrics

We now examine how generic properties of instances affect the quality of decisions and time-on-task. We study two types of properties: feature-space and solution-space metrics. The main difference between them is that feature-space metrics can be estimated from mathematical properties of the instance without any knowledge of an instance’s solution, whereas the calculation of solution-space metrics require knowledge of an instance’s solution, that is, require the solution to be computed (Fig 1). This makes estimating solution-space metrics more computationally (and epistemologically) intensive.

We first examine the effect of typical-case complexity (TCC), a feature-space metric of complexity, on human performance. This measure is based on a framework in computer science developed to study the drivers of computational hardness of computational problems by analyzing the difficulty of randomly generated instances of those problems. The study of random instances has revealed that there is substantial variation in computational resource requirements of instances with the same input length (*13*–*15*, *18*). This variation in computational hardness has recently been related to various structural properties of instances. In particular, it has been shown for several intractable (specifically, NP-complete) problems, including the KP (*18*), TSP (*13*) and 3SAT (*16*, *17*), that there exists a set of parameters 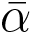 that captures the constrainedness of an instance. Moreover, it has been shown that there is a threshold *α*_*s*_ such that random instances with *α ≪ α*_*s*_ are mostly satisfiable while those with *α* ≫ *α*_*s*_ are mostly unsatisfiable. Importantly for our study, it has been shown for each of the problems under consideration that algorithms, on average, require more computational resources (e.g., time) to solve instances near *α*_*s*_ (*13*–*15*, *18*). In our study, we sampled instances with varying values of *α* and categorize instances with *α* ~ *α*_*s*_ as instances with a *high TCC* and instances with *α* ≫ *α*_*s*_ or *α ≪ α*_*s*_ as instances with *low TCC* (see Fig 2 and Materials and Methods).

**Figure 2:**
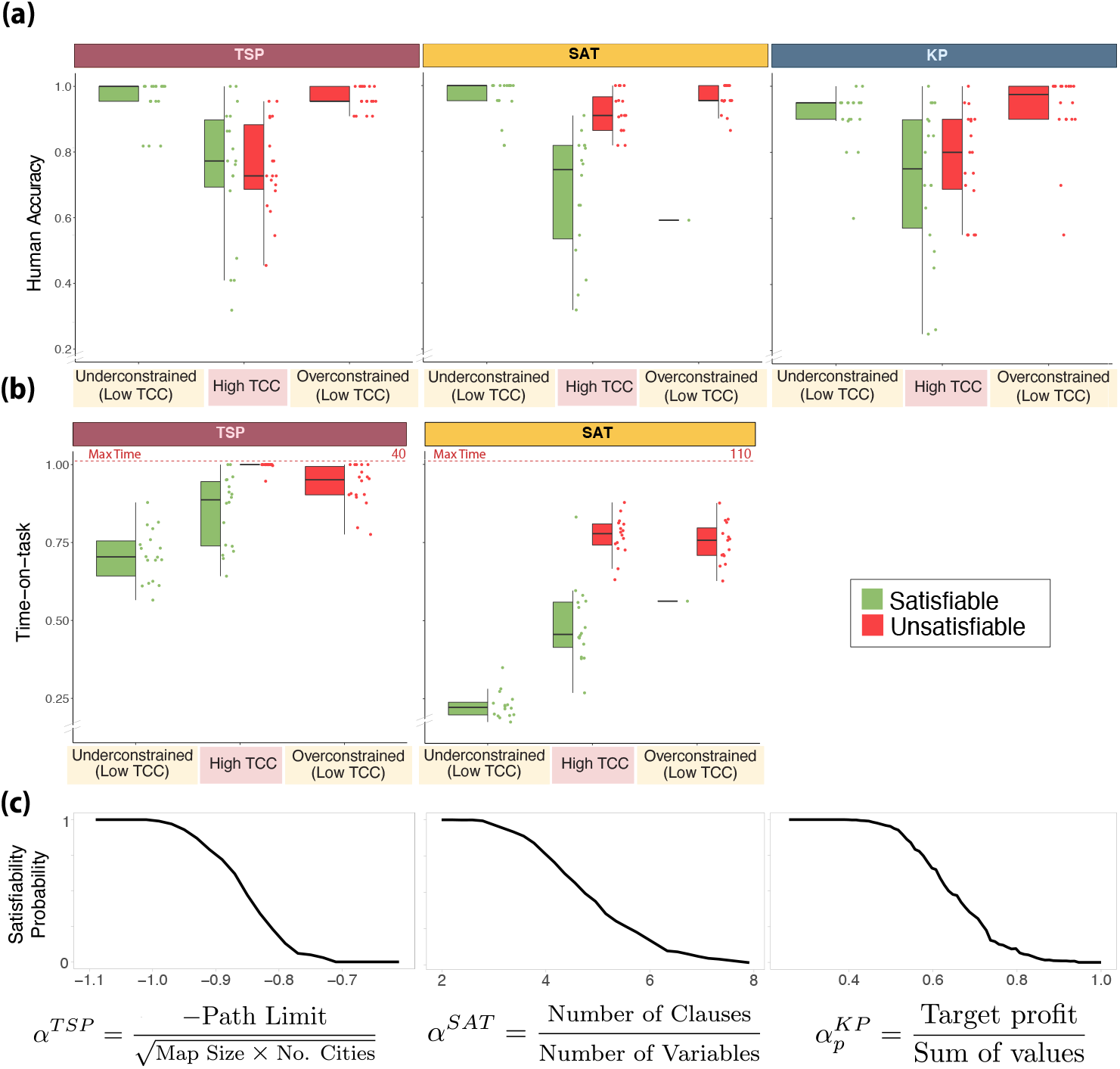
Typical-case complexity (TCC). **(a) Human accuracy.** Each dot represents an instance of one of the three problems considered. For each instance, human accuracy corresponds to the proportion of participants that solved an instance correctly. The instances are categorized according to their constrainedness region (*α*) and their TCC. The correct solution (satisfiability) of an instance is represented by its color. **(b) Time-on-task.** Median time spent solving an instance before submitting an answer. Time is represented as a proportion of the maximum time allotted on each trial (40s in the TSP and 110s in the 3SAT). **(c) Satisfiability probability and constrainedness parameter** *α*. Probability that a random instance is satisfiable as a function of *α* (the probability is empirically estimated; see Materials and Methods). In the underconstrained region (low TCC) the satisfiability probability is close to one while in the overconstrained region (low TCC) the probability is close to zero. The region with a high TCC corresponds to a region in which the probability is close to 0.5. *The box-plots represent the median, the interquartile range (IQR) and the whiskers extend to a maximum length of 1.5*IQR*.

We hypothesized that participants would have lower accuracy on instances with high TCC compared to those with low TCC. We found that this was indeed the case for both TSP and 3SAT and consistent with earlier results for KP (TSP: *β*_0.5_ = −2.10, *HDI*_0.95_ = [−2.50, −1.73], Table S2 Model 2; 3SAT: *β*_0.5_ = −1.58, *HDI*_0.95_ = [−1.95, −1.20], Table S1 Model 2; KP: *β* = −1.327 *P* < 0.001, main effect of TCC on accuracy, GLMM; Fig 2a).

Instances with *α* ≫ *α*_*s*_ or *α* ≪ *α*_*s*_ are considered to have low TCC. However, these instances belong to two structurally different regions, namely an overconstrained and an underconstrained region. We then studied whether differences in constrainedness affected accuracy among low TCC instances. We found that for the TSP and 3SAT, there was no difference in accuracy between underconstrained and overconstrained regions (TSP: *β*_0.5_ = 0.14, *HDI*_0.95_ = [−0.58, 0.87], Table S2 Model 3; 3SAT: *β*_0.5_ = −0.43, *HDI*_0.95_ = [−1.12, 0.28], Table S1 Model 3; the difference in effect, *overconstrained* − *underconstrained*, on accuracy, GLMM). These results are consistent with those obtained previously in relation to KP (*β* = 0.250, *P* = 0.355, the difference in effect, *overconstrained* − *underconstrained*, on accuracy, GLMM; Fig 2a). Taken together, these findings suggest that the mapping between *α* and TCC captures the effect of *α* on accuracy.

We also expected TCC to have an effect on time-on-task. We hypothesized that participants would spend more time on instances with high TCC. We found this to be the case for 3SAT and TSP (3SAT: *β*_0.5_ = 0.149, *HDI*_0.95_ = [0.116, 0.182], Table S3 Model 2; TSP: *β*_0.5_ = 0.118, *HDI*_0.95_ = [0.090, 0.147], Table S4 Model 2; effect of TCC on time-on-task as a proportion of the maximum possible time, censored linear mixed-effects models (CLMM), Fig 2b).

Furthermore, we explored whether constrainedness (*α*) had an effect on time-on-task, beyond the one captured by TCC. We found that participants spent less time-on-task on instances in the underconstrained region (3SAT: *β*_0.5_ = −0.352, *HDI*_0.95_ = [−0.385, −0.318] Table S3 Model 3; TSP: *β*_0.5_ = −0.199, *HDI*_0.95_ = [−0.233, −0.164], Table S4 Model 3; difference in time-on-task between instances in the underconstrained region and those with *α* ~ *α*_*s*_ (high TCC), CLMM). In the TSP, participants spent less time on overconstrained instances compared to those instances with *α* ~ *α*_*s*_, but this effect was not significant (*β*_0.5_ = −0.024, *HDI*_0.95_ = [−0.059, 0.011], difference in time-on-task between instances in the overconstrained region and *α* ~ *α*_*s*_, CLMM; Table S4 Model 3). In contrast, in the 3SAT participants spent more time on overconstrained regions compared to those instances with *α* ~ *α*_*s*_ (*β*_0.5_ = 0.071, *HDI*_0.95_ = [0.036, 0.106], difference in time-on-task between instances in the overconstrained region and with high TCC, CLMM; Table S3 Model 3). It is worth noting that in 3SAT, the more constrained the problem is, the more clauses are presented and, therefore, the higher the amount of information that needs to be parsed. This suggests that time-on-task in SAT would be driven by both the number of clauses and TCC. In line with this, we found evidence of a linear effect of the number of clauses on time-on-task even when controlling for TCC (*β*_0.5_ = 0.02, *HDI*_0.95_ = [0.02, 0.02], Table S9 Model 2). Interestingly, the number of clauses did not have a significant effect on accuracy (*β*_0.5_ = −0.02, *HDI*_0.95_ = [−0.05, 0.01], Table S9 Model 2). It is worth highlighting that the effect of TCC was still significant, on both time and accuracy, when controlling for the number of clauses (Table S9).

Our results so far show that participants expend more time on instances with higher TCC and yet they perform worse on these instances. This suggests a negative correlation between time-on-task and accuracy, which we corroborated (TSP: *β*_0.5_ = −0.1, *HDI*_0.95_ = [−0.13, −0.08], Table S2 Model 5; 3SAT: *β*_0.5_ = −0.02, *HDI*_0.95_ = [−0.02, −0.01], Table S1 Model 5); effect of time-on-task on accuracy, GLMM).

### Solution-space complexity metrics

In the previous section, we studied the effects of constrainedness and TCC, feature-space metrics, on human performance. These metrics can be estimated based on a problem’s input, that is, without the need to solve the instance. However, they characterize complexity as an average over a collection of random instances. We now explore a collection of related metrics, which although more computationally expensive, are expected to provide a more detailed account of human performance. These metrics are based on an instance’s *solution space*. We will use the term solution space to refer to the set of *solution witnesses* of an instance, that is, the set of configurations of variables (e.g., possible paths or variable assignments) that satisfy an instance’s constraints. Note that in order to estimate solution-space metrics, the instance, or a harder variant, has to be solved. In some cases, all possible solution witnesses must be found.

#### Satisfiability

A relevant solution-space property of instances, is their *satisfiability*, that is, whether the instance’s solution is ‘yes’ or ‘no’. We found that satisfiability affects accuracy but that this effect varies between problems. In 3SAT, participants performed worse on satisfiable instances (*β*_0.5_ = −1.35, *HDI*_0.95_ = [−1.73, −0.99], main effect of satisfiability, GLMM, Table S1 Model 8), whereas there was no significant effect of satisfiability on accuracy in the TSP and the KP (TSP: *β*_0.5_ = −0.06, *HDI*_0.95_ = [−0.34, 0.22], Table S2 Model 6; KP: *β*_0.5_ = −0.29, *HDI*_0.95_ = [−0.57, 0.01], Table S6 Model 1; main effect of satisfiability, GLMM). Turning our attention to the effect of satisfiability on time-on-task, we find that less time was spent on satisfiable instances in both TSP and 3SAT (TSP: *β*_0.5_ = −0.17, *HDI*_0.95_ = [−0.20, −0.15], Table S4 Model 4; 3SAT: *β*_0.5_ = −0.32, *HDI*_0.95_ = [−0.35, −0.29], Table S3 Model 4; effect of satisfiability on time-on-task, CLMM). In summary, we observed that participants spent less time-on-task on satisfiable instances in both TSP and 3SAT, yet the effect of satisfiability on accuracy varied across problems.

We then gauged whether the effect of satisfiability on performance was modulated by TCC. Our results suggest that satisfiability and TCC interact and affect accuracy and time-on-task only in the 3SAT. (Fig 2; S1 Appendix).

#### Number of solution witnesses

We can analyze the drivers of hardness in satisfiable instances at a more granular level by studying the number of solution witnesses of an instance. This generic feature of decision instances captures the constrainedness of an instance: a higher value of witnesses is related to a lower degree of constrainedness. This metric, however, is only informative for satisfiable instances (by definition, unsatisfiable instances have zero solution witnesses). Thus, we restrict our analysis to these instances.

We found, in line with our hypothesis, a positive effect of the number of witnesses on accuracy in all three problems (3SAT: *β*_0.5_ = 0.62, *HDI*_0.95_ = [0.49, 0.79]; TSP: *β*_0.5_ = 0.45, *HDI*_0.95_ = [0.37, 0.53]; KP: *β*_0.5_ = 0.26, *HDI*_0.95_ = [0.19, 0.34]; main effect of the number of witnesses on accuracy, GLMM; Table S5 Models 1,4,6; Fig 3a). We further hypothesized that participants would spend less time solving instances with a higher number of witnesses. This was indeed the case (TSP: *β*_0.5_ = −0.02, *HDI*_0.95_ = [−0.03, −0.02], Table S4 Model 5; 3SAT: *β*_0.5_ = −0.041, *HDI*_0.95_ = [−0.047, −0.036], Table S3 Model 5; effect of number of witnesses on time-on-task, CLMM; Fig 3b)). These results suggests that, among satisfiable instances, the more constrained an instance, the harder it is to solve.

**Figure 3:**
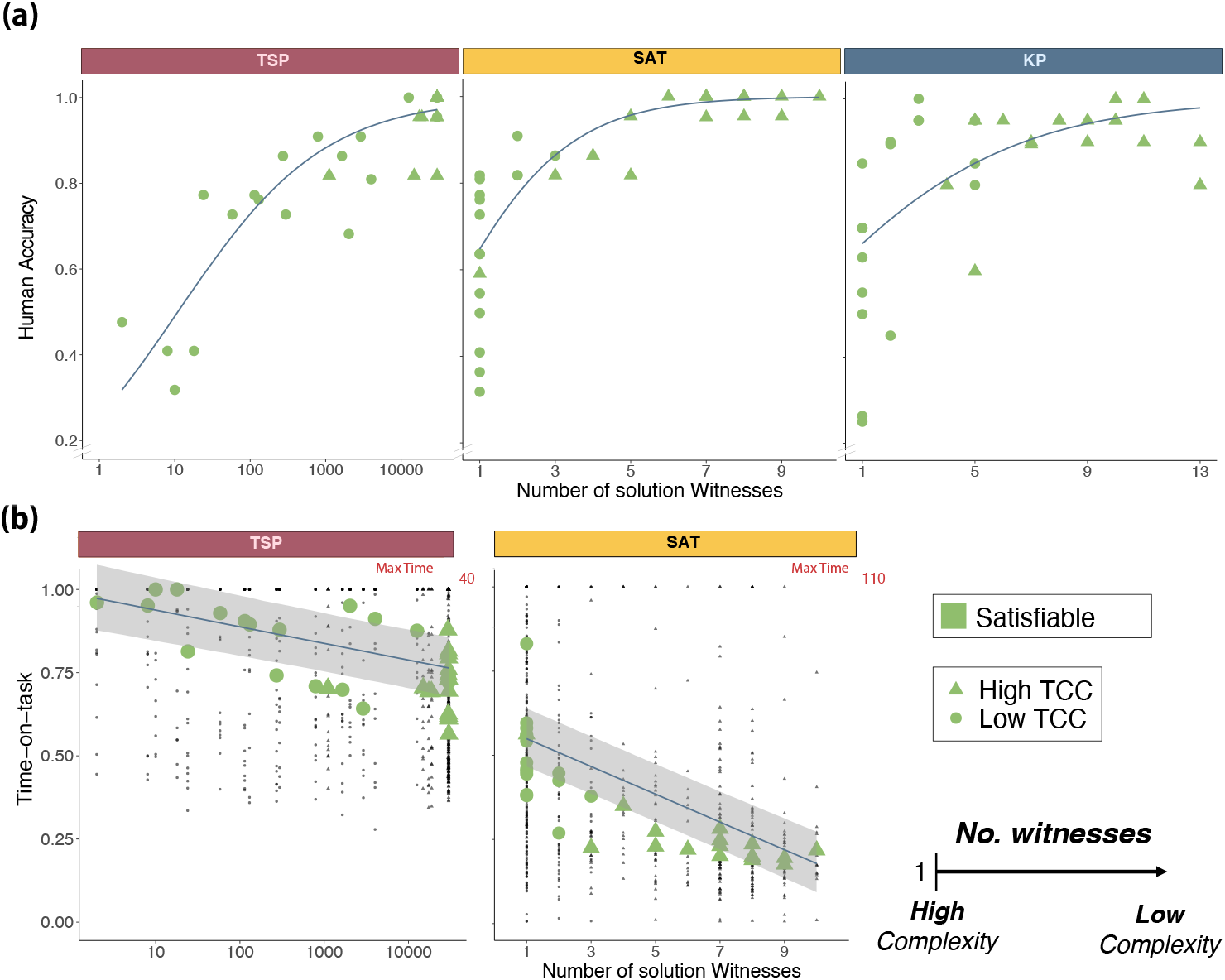
Number of solution witnesses. The number of witnesses is defined as the number of *state-space combinations* (i.e., paths, selection of items or switch-setups) that satisfy the constraints. On satisfiable instances, the problem becomes harder as the number of witnesses approaches 1. Only satisfiable instances are included. **(a) Human accuracy.** Each green shape represents the mean accuracy per instance. The blue line represents the marginal effect of the number of solution witnesses on human accuracy (GLMM Table S5 Models 1,4,6). **(b) Time-on-task.** Each green shape represents the median time-on-task per instance. The blue line represents the marginal effect (and 95% credible interval) of the number of solution witnesses on time-on-task (LMM Table S4 Model 5 and Table S3 Model 5). Each black dot corresponds to the time-on-task of one participant while solving a single instance.

The number of witnesses is a metric conceptually similar to TCC. While TCC is a mapping from expected constrainedness (*α*) to hardness, the number of witnesses maps the realized level of constrainedness of a single instance to hardness. Therefore, we expected the number of witnesses to drive the effect of TCC on performance in satisfiable instances. Our results, indeed, suggest that the effect of TCC on accuracy in satisfiable instances is driven, at least partially, by the number of witnesses of an instance (see S4 Appendix).

#### Instance complexity

Although the number of solution witnesses captures invariants of human behavior in all three problems, it does so only for satisfiable instances. We therefore now explore an alternative solution-space metric, instance complexity (IC), that can be used to study the difficulty of all instances (both satisfiable and unsatisfiable) (*22*). IC is related to the constrainedness of an instance and the order parameter 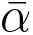. It is characterized based on the distance between the decision threshold of an instance and the optimal attainable value in the optimization variant of the instance. For example, the optimization variant of an instance of the TSP corresponds to finding the minimum path-length connecting all cities. For the KP, it corresponds to finding the maximum value that can fit into the knapsack given the weight constraint and given the same set of items from the decision variant. Analogously, for the 3SAT, the optimization version (MAX-SAT) corresponds to finding the maximum number of clauses that can be rendered true simultaneously.

We then define IC as the absolute value of the normalized difference between target value of the decision variant and the optimal value attainable of the corresponding optimization variant. In the KP, for example, it is defined as follows,

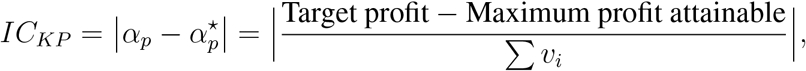

where Σ*v*_*i*_ is the sum of values of all the items. In other words, in the KP, IC is the normalized value of the distance between the target profit of a decision instance and the maximum profit that can be attained with the same set of items and the same capacity constraint. The corresponding expressions for TSP and 3SAT are provided in the Methods section.

We studied the effect of IC on performance in each of the problems. Note that lower values of IC indicate that the decision threshold is closer to the optimum, which corresponds to a higher level of computational hardness. Therefore, we expected a positive relation between IC and accuracy. We found a positive non-linear relation in all problems (KP: *β*_0.5_ = 9.05, *HDI*_0.95_ = [7.20, 11.02], Table S6 Model 2; TSP: *β*_0.5_ = 21.13, *HDI*_0.95_ = [17.63, 24.91], Table S2 Model 7; 3SAT: *β*_0.5_ = 30.30, *HDI*_0.95_ = [21.95, 39.24], Table S1 Model 6; the effect of *IC* on accuracy, GLMM; Fig 4a).

**Figure 4:**
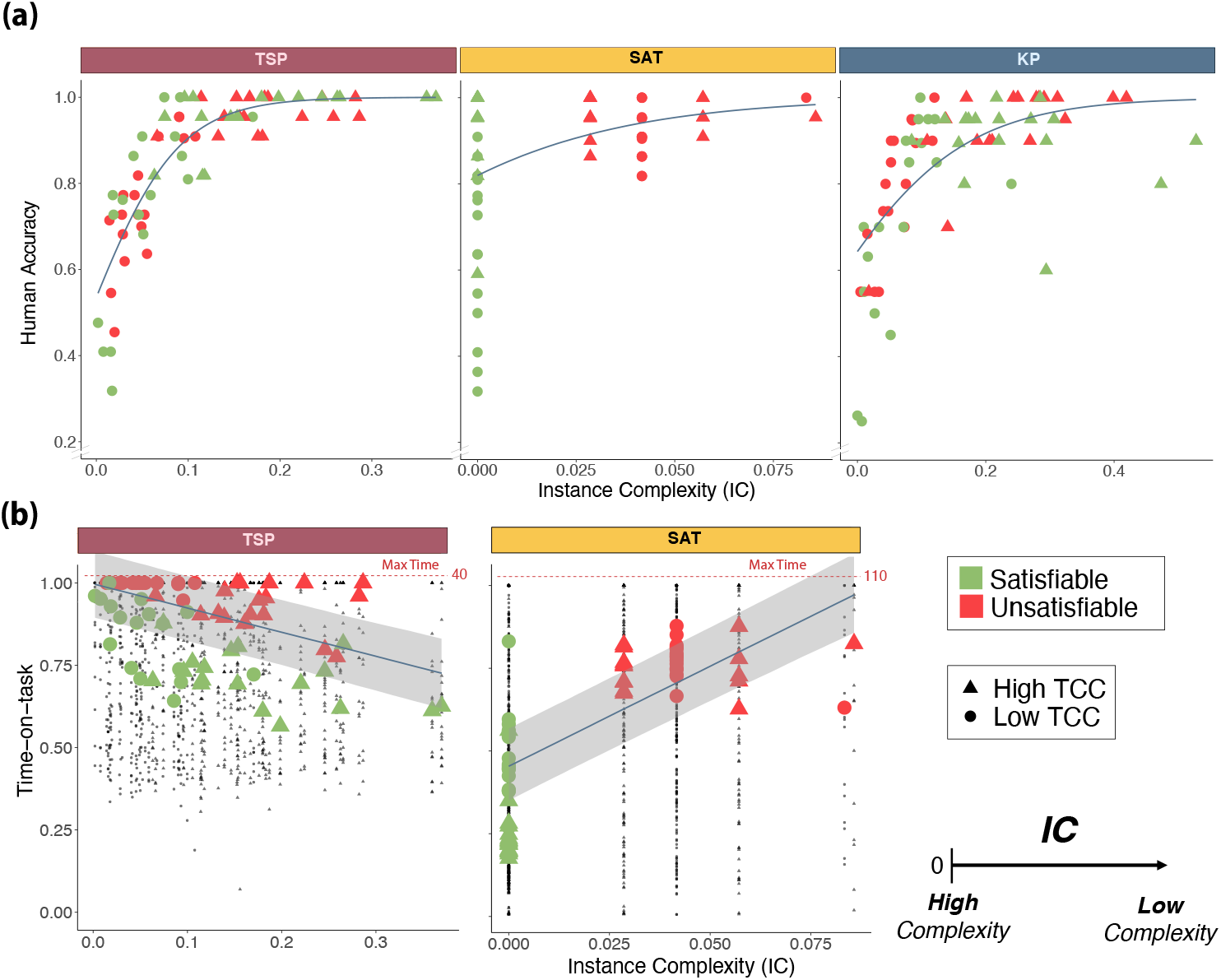
Instance complexity. Instances become harder as 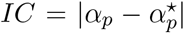 approaches 0. **(a) Human accuracy.** Green and orange shapes represent the mean accuracy for each instance. The blue lines represents the marginal effect of IC on human accuracy (GLMM Table S6 Model 2, Table S2 Model 7, Table S1 Model 6). **(b) Time-on-task.** Green and orange shapes represent the median time-on-task for each instance of the TSP and 3SAT problems. The blue lines represents the marginal effect (and 95% credible interval) of IC on time-on-task (LMM Table S4 Model 6, Table S3 Model 6). Each black dot corresponds to the time spent by a single participant on a particular instance.

IC is a metric that overcomes previous limitations of TCC, in that it is a metric of hardness of individual instances. Moreover, it provides information beyond the number of solution witnesses since it captures variability across unsatisfiable instances as well. As such, we expected IC to capture a substantial amount of the variability in human accuracy between instances. Indeed, *IC* was able to explain a high proportion of the variance in average instance accuracy in the TSP and the KP (KP: *R*^2^ = 0.47; TSP: *R*^2^ = 0.74; fraction of variance explained by Table S6 Model 2 and Table S2 Model 7; the target variable to explain is the mean accuracy per instance), but was lower in 3SAT (3SAT: *R*^2^ = 0.17; fraction of variance explained by Table S1 Model 6). These results indicate that IC is able to explain variance in accuracy across tasks, but to a lesser degree in 3SAT. This is even after considering the characteristics of the sampled 3SAT instances in which this is tested. Indeed, the effect of IC on accuracy in the 3SAT might be incongruously driven by satisfiability (S5 Appendix).

Next, we explored the effect of IC on time-on-task. We expected a negative relation between IC and the average time spent on an instance. This was the case for TSP (*β*_0.5_ = −0.735, *HDI*_0.95_ = [−0.901, −0.581], main effect of IC on time-on-task, CLMM; Table S4 Model 6; Fig 4b), but for the 3SAT we found a significant positive effect (*β*_0.5_ = 6.04, *HDI*_0.95_ = [5.41, 6.70], main effect of IC on time-on-task, CLMM; Table S3 Model 6; Fig 4b). Based on this result, we hypothesized that the positive effect of IC on time-on-task in 3SAT could have been driven by the effect of satisfiability, but we are unable to test this hypothesis directly. Therefore, we investigated the effect of IC on time-on-task in unsatisfiable instances only and found a non-significant negative effect (*β*_0.5_ = −0.557, *HDI*_0.95_ = [−1.912, 0.782], main effect of IC on time-on-task for unsatisfiable instances, CLMM; Table S3 Model 7). These results indicate a negative relation between IC and time-on-task in the TSP, whereas in 3SAT the results are inconclusive.

Finally, we validate our results in an alternative measure of performance. We have shown that generic instance-level complexity metrics are able to explain differences in two dimensions of performance: accuracy and time-on-task. The latter, in particular, is associated with the level of cognitive effort exerted. In order to validate whether the effect of the proposed complexity metrics on time-on-task could be extended to other aspects of cognitive effort, we explored the effect of these metrics on the number of clicks participants performed in each trial. Arguably, participants search the state space by clicking on different state combinations in order to decide whether an instance is satisfiable or not. Differences in the quantity of clicks used to solve an instance can shed light into the length of search. We find that the effect of generic properties of instances on time-on-task is qualitatively replicated on the number of clicks, that is, participants used more clicks on harder instances. Search was longer in general in the case of unsatisfiable instances and there was a positive effect of TCC on search length. Moreover, longer search was also related to lower values of IC and lower number of witnesses (see S2 Appendix).

## Discussion

Computational complexity theory has been used to study the limits of what is potentially human computable (*2*, *4*, *25*). According to this work, many tasks we face in our lives—and corresponding computational models of human behavior—are computationally infeasible, including planning, learning and many forms of reasoning (for example, analogy, abduction and Bayesian inference) (*4*). However, this analysis is not suited to, and does not aim to, explain differences in performance and behavior across the class of human-computable problems. Such differences have generally been ascribed to the solver or the agent (*9*–*12*, *26*–*28*). This approach, however, is difficult to generalize given the diversity of algorithms implemented and their specificity to particular problems (*10*, *29*).

We propose that computational complexity theory can inform the study of cognitive capacities beyond studying the a priori plausibility of cognitive models. Specifically, we put forward that this theory can be used to study human performance. In the present study, we show that this conceptual approach captures differences in behavior across different NP-complete problems without reference to an algorithm or particular computational device. This generic framework quantifies computational hardness of cognitive tasks based on structural properties of individual instances of the the underlying computational problem. The results of this study show that a set of metrics based on these properties predict both task accuracy and time-on-task across three cognitive tasks related to different NP-complete computational problems.

More specifically, using a controlled experiment, we show that three generic properties of NP-complete problems, typical-case complexity (TCC), the number of solution witnesses, and instance complexity (IC), affect human performance when performing a task. While the extent of time-on-task increased with higher complexity of instances, accuracy, and thus efficacy in those instances, decreased. We show that the relation between the complexity metrics presented on the one hand and task performance on the other, are similar across three different types of NP-complete problems. This is particularly surprising because of the diverse nature of these problems. While the SAT task is an abstract logical problem, TSP is presented as a visual navigation task and the KP is associated with arithmetic calculations.

Our results complement findings from computer science and suggest that hardness stems partially from the intrinsic difficulty of the problem and the instance, regardless of the algorithm and the computing device used. In particular, our findings suggest that the same intrinsic hardness metrics describe performance of algorithms executed by both electronic computers (*13*–*18*) and humans, that is, biological computers. This is particularly interesting because the theory in which our analysis is based is derived without taking into account limits on human computation. For instance, no memory constraints are imposed on the solving algorithms. Interestingly, our results also show that computational hardness affects how much time an agent decides to spend on an instance. This is far from obvious because, unlike standard algorithms executed by electronic computers, humans have the option to stop working independently of the solving strategy.

### Metrics of complexity

We explored the effect of three generic complexity metrics on human performance. Importantly, while the proposed metrics capture the computational hardness of NP-complete problems, they can be applied to other decision problems in classes P or NP (*14*–*16*), and have been shown to be extendable to optimization problems (*22*). Each of the metrics can be used to unearth generalities in human behavior. Typical-case complexity (TCC) captures the average hardness of a random ensemble of instances of a problem based on its constrainedness. Critically, TCC can be computed ex-ante—without knowledge of an instance’s solution. The number of solution witnesses captures a structural property of a single instance that is related to the hardness of search for satisfiable instances. Finally, instance complexity (IC) maps constrainedness of a single instance to computational hardness, for both satisfiable and unsatisfiable instances.

The generality of TCC is limited by its dependence on a particular sampling distribution. We sampled instances for each of the problems from a specific procedure in which the components of the instances were randomly sampled from uniform distributions. We leave it to future research to study whether TCC can be extended to other probability distributions, and particularly, to those found in real life (*30*).

Importantly, we provided two alternatives to TCC (IC and number of witnesses), which do not depend on a sampling procedure. These metrics quantify the hardness of specific instances of problems. However, they do come at a cost: these metrics are computationally intensive. That is, in order to compute them, the decision problem, or a harder variant, needs to be solved first. For IC to be estimated, the optimization variant of the instance needs to be solved, whereas to compute the number of witnesses, all of the possible witnesses of an instance need to be counted.

We argue that the computational requirements of calculating these metrics is not prohibitive in the context of the study of human problem-solving and cognition in general. These metrics can be used to predict generalities in human behavior with the aid of any of the resources at hand, including electronic computers. Therefore, since the practical instances of problems solvable by humans are relatively small compared to those solvable by electronic computers, cognitive scientists effectively have access to an oracle machine to estimate computationally intensive metrics.

### Future directions

This paper focuses on generalities across problems within a well-defined class (i.e., NP-complete). A related question is whether intrinsic characteristics specific to a problem can complement the generic metrics presented here. Intrinsic metrics of complexity, specific to a problem, have been previously shown to affect performance. Specifically, for all three problems considered in this study, measures derived from the features of the problem have been shown to affect computational time of algorithms executed on electronic computers (*31*–*34*). Additionally, problem-specific complexity metrics have been show to be related to human performance in the optimization variants of the TSP (*6*, *7*). Future work should be undertaken to study how task-independent and problem-specific metrics jointly affect human performance.

Importantly, our results suggest that the metrics put forward in this study are generic as they provide both ex-ante (i.e., before solving the instance) and ex-post (i.e., after solving the instance) predictability across different problems. However, our work also highlights that certain structural properties of the problem might have problem-specific effects that could interact with the effect of generic metrics of hardness. This is particularly evident with satisfiabilty, which has a differential effect across problems. Explicitly, in the 3SAT task, unlike in the other two, satisfiability has a significant effect on accuracy. Indeed, the effect of IC on accuracy in the 3SAT might be incongruously driven by satisfiability, in a way that cannot be differentiated with our experimental design (all 3SAT satisfiable instances have *IC* = 0). This could partially explain why we find that IC explains less of the variance in accuracy in the 3SAT than in the other two tasks and that the effect of IC on time-on-task is inconclusive. Future studies should attempt to disentangle these effects, for example, by studying the related maximum satisfiability problem (MAX-SAT). More importantly, these results warrant further investigation of the effect of satisfiability, and other structural properties, on human behavior. Moreover, future work could explore the differences in these effects across classes of problems. For instance, NP-complete problems could be categorized into finer classes based on the effect of particular properties on human problem-solving. Our findings would suggest that more abstract logical problems might be solved differently to other more life-pertinent problems, such as KP and TSP.

A related open question is how the nature of the presentation of the problem to the agent affects complexity. One conjecture would be that for humans, visual presentation may affect results only in absolute terms (e.g., one representation leads to higher average performance), but not in relative terms (i.e., relative difficulty between instances). Future research should determine whether presentation matters, in analogy with the present study, which shows that the nature of the NP-Complete problem does not matter.

We investigated the effect of different metrics of instance-level complexity keeping the size of instances fixed. An additional dimension in this framework that has been shown to affect human behavior is an instance’s size (*6*, *11*) and the size of the state-space (that is, the number of possible combinations or paths) (*10*, *35*). Additionally, the instance complexity metrics we presented are based on the satisfiability threshold and the number of witnesses. Recently, it has been shown that the performance of algorithms, designed for electronic computers, as *α* approaches *α*_*s*_, is not only related to the decrease in the number of witnesses, but also to the shattering of witnesses into distinct clusters (*20*, *36*). Further research is needed to integrate these different dimensions of complexity and determine their combined effect on human problem-solving.

We have argued that the framework presented here can be used to characterize the effect of the task environment on human performance. However, instance-level complexity metrics can also shed light on the type of strategies employed by agents. Note, for example, that our results suggest that participants did not predominantly perform random search. In a random algorithm, random combinations from the *state-space* (i.e., paths or variable configurations) are tried and an answer (yes/no) is selected depending on whether a solution witness is found. If participants implemented such an algorithm, we would expect the length of search to be similar on all unsatisfiable instances given that completing an exhaustive search of the state space is unlikely because of the limits on time and number of clicks. This, however, is not what we found. In unsatisfiable instances the length of search, time-on-task (in TSP) and accuracy were affected by IC. In fact, by applying the same argument, we can rule out other more directed search heuristics such as greedy algorithms (*37*), which have been proposed to be linked to human behavior (*10*). Overall, our results suggest that when people solve decision problems, they implement procedures to exclude alternatives from the witness solution set. If this was not the case, we would not find any effect of our complexity metrics among unsatisfiable instances. Further research is needed in order to disentangle between prospective algorithms and how this framework could be used to inform the study of strategy-use in humans.

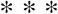

We provide empirical evidence that studying the intrinsic computational hardness of the task environment predicts human performance in a task. This approach allows for the objective characterization of what it means for an instance to be more difficult than another and the linking of these specifications to levels of performance. This principled approach may provide policy prescriptions (about when, e.g., financial products become excessively complex) that do not require one to take a stance on the strategies people employ (e.g., default options in retirement options rely on people choosing the first salient option). Indeed, in cases where the cognitive demands of a task substantially exceed decision-makers’ capacities, there is a need to prevent harm. One way to do so could be AI-powered applications that support people in making complex decisions. Another approach could involve regulatory interventions imposing limits on the complexity of products and/or require product and service providers to generate mechanisms to overcome complexity in the cases where a task is computationally hard and an agent’s cognitive capabilities are not sufficient to guarantee a high-quality decision. However, before such mechanisms can be developed and implemented, a well-founded characterization of the complexity of these tasks is needed. Here we present a milestone towards such a characterization.

## Materials and Methods

### Ethics statement

The experimental protocol was approved by the University of Melbourne Human Research Ethics Committee (Ethics ID 1749594.2). Written informed consent was obtained from all participants prior to commencement of the experimental sessions. Experiments were performed in accordance with all relevant guidelines and regulations, including the Declaration of Helsinki.

### Participants

A total of 47 participants were recruited in two separate groups from the general population (group 1: 24 participants; 12 female, 12 male; age range = 19-35 years, mean age = 24.1 years; group 2: 23 participants; 13 female, 10 male; age range = 18-32 years, mean age = 23.3 years). Inclusion criteria were based on age (minimum=18 years, maximum=35 years) and normal or corrected-to-normal vision.

Each group of participants were asked to solve a set of random instances of a computational problem. Group 1 participants were presented with 64 instances of the Boolean satisfiability problem (3SAT). Group 2 participants were presented with 72 instances of the decision variant of the traveling salesperson problem (TSP). Some trials and participants were excluded due to different issues (see Statistical Analysis in Materials and Methods).

### Experimental tasks

#### Boolean satisfiability task

This task is based on the Boolean satisfiability problem (3SAT). In this problem, the aim is to determine whether a Boolean formula is *satisfiable*. In other words, given a propositional formula, the aim is to determine whether there exists at least one configuration of variables (which can take values TRUE or FALSE) such that the formula evaluates to TRUE. The propositional formula in 3SAT has a specific structure. Specifically, the formula is composed of a conjunction of clauses that must all evaluate TRUE for the whole formula to evaluate TRUE. Each of these clauses, takes the form of an OR logical operator of three literals (variables and their negations). An example of a 3SAT problem is:

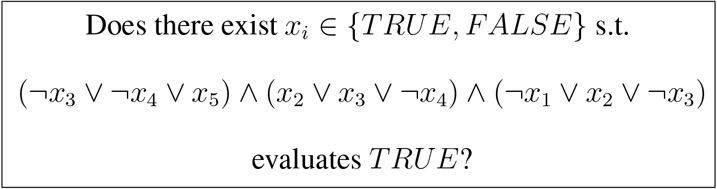

In order to represent this in an accessible way to participants we developed a task composed of switches and light bulbs (Fig 5a). Participants were presented with a set of light bulbs (clauses), each of which had three switches (literals) underneath that were represented by a positive or negative number. The number on each switch represented the variable number, which could be turned on or off (TRUE or FALSE). The aim of the task is to determine whether there exists a way of turning on and off variables such that all the light bulbs are turned on (that is, the formula evaluates TRUE).

**Figure 5:**
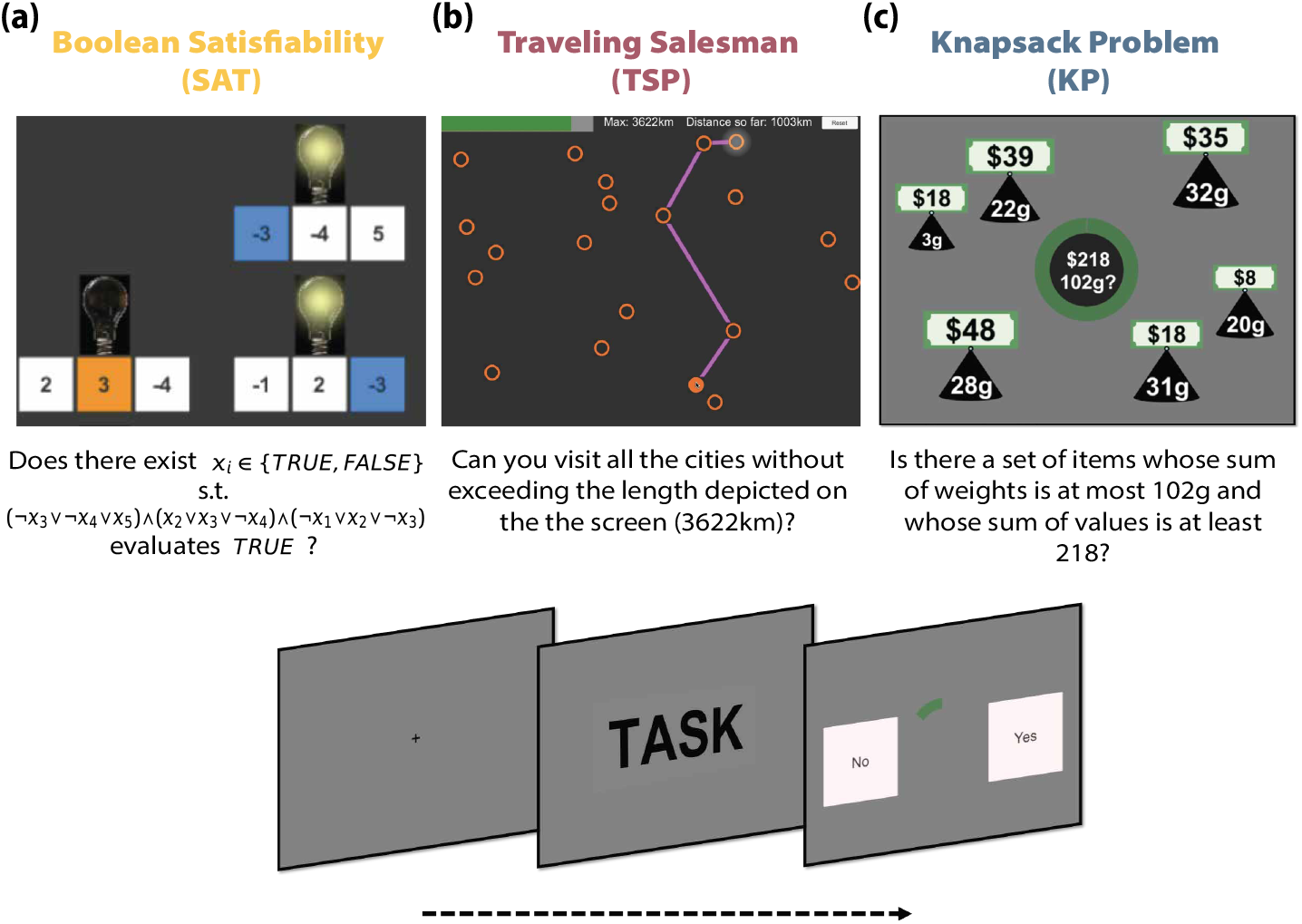
Experimental tasks. **(a) Boolean satisfiability.** In this task, the aim is to determine whether a Boolean formula is *satisfiable*. The Boolean formula is represented with a set of light bulbs (clauses), each of which has three switches (literals) underneath that are characterized by a positive or negative number. The number on each switch represents the variable number, which can be turned on or off (TRUE or FALSE). The aim of the task is to determine whether there exists a way of turning on and off variables such that all the light bulbs are turned on (the corresponding Boolean formula evaluates to TRUE). The task was interactive. Participants could click on switches to turn them on and the corresponding literals and light bulbs would change color automatically. This stage had a time limit of 110 seconds. Afterwards, participants had 3 seconds to make their response (either a ‘YES’ or ‘NO’). **(b) Traveling salesperson.** Participants are given a list of cities displayed on a rectangular map on the screen and a limit *L* on path length. The problem is to determine whether there *exists* a path connecting all *N* cities with a distance at most *L*. The task was interactive. Participants could click from city to city and the corresponding path and distance traveled would display and update automatically. This stage lasted a maximum of 40 seconds. Afterwards, participants had 3 seconds to make their response. **(c) Knapsack.** Participants are presented with a set of items with different values and weights. Additionally, a capacity constraint and target profit are shown at the center of the screen. The aim is to ascertain whether there exists a subset of items for which (1) the sum of weights is lower or equal to the capacity constraint and (2) the sum of values yields at least the target profit. The task was not interactive. This stage lasted for 25 seconds. Finally, participants had 2 seconds to make their response.

At the beginning of each trial, participants were presented with a different instance of the 3SAT problem. A bar in the top-right corner of the screen indicated the time remaining in the trial. Each participant completed 64 trials (4 blocks of 16 trials with a rest period of 60 seconds between blocks). Trials were self-paced with a time limit of 110 seconds. Participants could use the mouse to click on any of the variables to select their value ({*blue* = *TRUE, orange* = *FALSE*}). A light bulb above each clause indicated whether a clause evaluated to TRUE (light on) given the selected values of the variables underneath it. The number of clicks in each trial was limited to 20. The purpose of this limit was to discourage participants from using a trial-and-error strategy to solve the instances. When participants were ready to submit their solution, they pressed a button to advance from the screen displaying the instance to the response screen where they responded YES or NO. The time limit to respond was 3 seconds, and the inter-trial interval was 3 seconds as well. The order of instances and the side of the YES/NO button on the response screen were randomized for each participant.

##### Instance sampling

A random instance is a selection of clauses and literals in which *M* clauses of three literals are chosen randomly. Each of the literals is associated with one of *N* variables. Both numerical (*16*) and analytical (*17*) evidence suggests that in the limit *N* → ∞, there exists a value of the clause to variables ratio *α* = *M/N*, 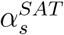, such that typical instances are satisfiable for 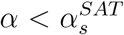, while typical instances are unsatisfiable for 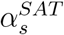. The current best estimate for the satisfiability threshold, 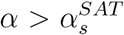, as *N* → ∞ is 4.267 (*17*) (note that the value of *α*_*s*_ is a function of the number of literals per clause, which was fixed at 3 in this study). As *N* → ∞, instances near the threshold are on average harder to solve (*14*, *15*). We exploit both the threshold phenomenon in satisfiability and its link to computational hardness.

We generated random instances with different degrees of complexity by varying *α*. We picked a value of *α*, starting at the lower bound of its range and incrementing in steps of 0.1 until the upper bound was reached. For each value of *α*, we computed the number of clauses *M* by multiplying *α* and a fixed value of *N* and then rounding to the nearest integer. *N* (and the time limit for the task) was determined before hand using pilot data to ensure that the task was not too easy nor too hard for participants (that is, to ensure sufficient variation in accuracy). Importantly, *N* was also restricted to values in which the corresponding number of clauses could fit in the screen of the task. Specifically, we restricted the number of clauses to be at most 36.

Once *N* was fixed, we generated 1000 random instances for each value of *M*. Each random instance was generated by first selecting the literals for each clause. Each literal is represented by a positive or negative sign (negation of a variable) and is sampled from the set {−1, +1} with equal probability. Afterwards, three variables were selected for each clause by sampling without replacement from the set of *N* variables (*14*, *16*).

From the randomly generated instances we first determined the satisfiability threshold of our finite instances (*N* = 5). That is, we calculated the value of 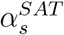 at which half of the randomly generated instances were satisfiable and half were unsatisfiable. This was the case for *α* = 4.8. Based on this we selected a subset of random instances to use in the task.

We asked participants to solve a set of instances randomly sampled from three different regions: an underconstrained region 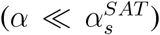, a region around the satisfiability threshold 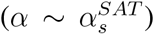 and an overconstrained region 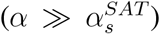. Instances near the satisfiability threshold are defined to have a *high TCC*, whereas instances further away from the satisfiability threshold (in the under-constrained or over-constrained regions) are defined to have a *low TCC*. We selected 16 instances from the underconstrained region (*α* = 2) and 16 instances from the overconstrained region (*α* = 7). We then sampled 32 instances near the satisfiability threshold (*α* = 4.8), such that 16 of the selected instances were satisfiable and 16 were not satisfiable.

In order to also ensure a sufficient degree of variability between instances near the satisfiability threshold, we added an additional constraint in the sampling procedure. For each set of instances (satisfiable and not satisfiable) we forced half to have algorithmic complexity less than the median algorithmic complexity at this value of *α*, and the other half to be harder than the median. The algorithmic complexity was estimated using an algorithm-specific complexity measure of a widely-used algorithm (*Gecode* propagations parameter). Gecode is a generic solver for constraint satisfaction problems that uses a constraint propagation technique with different search methods, such as branch-and-bound. We chose an output variable, the number of propagations, that indicates the difficulty for the algorithm of finding a solution and whose value is highly correlated with computational time. We did not use compute time directly as a measure of algorithmic complexity because for instances of small size, like the ones used in this study, compute time is highly confounded with overhead time. Thus, our set of instances in the region *α* ~ *α*_*s*_ comprised 8 instances in each of the following categories {satisfiable, unsatisfiable} × {low/high algorithmic difficulty}.

#### Traveling salesperson task

This task is based on the traveling salesperson problem. Given a set of *N* cities displayed on a rectangular map on the screen and a limit *L* on path length, the decision problem is to answer whether there *exists* a path connecting all *N* cities with a distance of at most *L* (Fig 5b).

In the TSP task, each participant completed 72 trials (3 blocks of 24 trials with a rest period of 30 seconds between blocks). Each trial presented a different instance of TSP. Trials were self-paced with a time limit of 40 seconds. Participants could use the mouse to trace routes by clicking on the dots indicating the different cities. The length of the selected route at each point in time was indicated at the top of the screen (together with the maximum route length of the instance). When participants were ready to submit their answer, they pressed a button to advance from the screen displaying the cities to the response screen where they responded YES or NO. The time limit to respond was 3 seconds, and the inter-trial interval was 3 seconds as well. The order of instances and the sides of the YES/NO buttons on the response screen were randomized for each participant.

##### Instance sampling

A TSP instance can be characterized as a collection of *N* cities, a matrix of distances ***d*** between each pair of cities, and a limit *L* on path length. Here, we restrict the problem to the euclidean TSP; that is, we constraint our distance matrices ***d*** to those that can be represented in a two-dimensional map of area *M*^2^.

Just like for 3SAT, it has been proposed that there exists a parameter *α*^*TSP*^ that captures the constrainedness of the problem, specifically suggests that in the 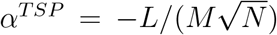 (*13*). Evidence limit *N* → ∞, there exists a value of *α*, 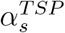, such that typical instances are satisfiable for 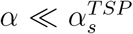, while typical instances are unsatisfiable for 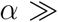 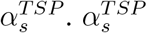 for the euclidean TSP is estimated at −0.7124 ± 0.0002 in the limit *N* → ∞ (*13*, *38*). As *N* → ∞ instances near 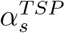 have been shown to be, on average, harder to solve (*13*). We use the this insight to vary typical-case complexity of instances of finite size.

Instances of the TSP had *N* = 20 cities. This value, and the time limit for the task, were determined using pilot data to ensure that the task was not too easy nor too hard for participants (that is, to ensure sufficient variation in accuracy). Random instances of the euclidean TSP were then generated by choosing (x,y) coordinates for each of the *N* = 20 cities uniformly at random from a square with side length *M* = 1000 (*13*). We generated 100 sets of coordinates; that is, 100 distance matrices ***d***. For each distance matrix, we generated instances with different values of *L*. We did this by varying the value of *α*, which was incremented in the range [−0.25, −1.25] with step size 0.02.

To determine the location of the satisfiability threshold in our sample of random instances (with *N* = 20), we determined the value of *α* at which half of the randomly generated instances were satisfiable and half were unsatisfiable. The satisfiability threshold was located at 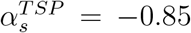. We randomly sampled instances at this value of *α* such that half of the selected instances were satisfiable and half were not satisfiable. We also ensured variability in the instances sampled near 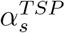 by forcing variability in the algorithmic complexity of sampled instances. Specifically, half of the instances had an algorithmic complexity (number of propagations) above the median and half below the median (see description of 3SAT above for details). Thus, our set of of instances in the region 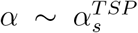 comprised 9 instances in each of the four following categories: {satisfiable, unsatisfiable} × {low/high algorithmic difficulty}.

For the underconstrained region, 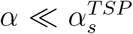, we randomly chose 18 instances from the set of 100 randomly generated instances with *α*^*TSP*^ = −0.99. For the overconstrained region, 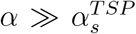, we randomly chose 18 instances from the set of 100 randomly generated instances with *α*^*TSP*^ = −0.71. We made sure that no two instances in our set of selected instances had the same set of city coordinates.

#### Knapsack task

In this paper we report on the experimental data collected on the knapsack decision task by Franco et al. (*22*). Statistical results from (*22*) were used when available.

The knapsack task is based on the 0-1 knapsack problem (KP). An instance of this problem consists of a set of items *I* = {1, … , *N*} with weights 〈*w*_1_, … , *w*_*N*_〉 and values 〈*v*_1_, … , *v*_*N*_〉, and two positive numbers *c* and *p* denoting the capacity and profit constraint (of the knapsack). The problem is to decide whether there exists a set *S* ⊆ *I* such that 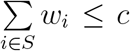, that is, the weight of the knapsack is less than or equal to the capacity constraint; and 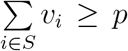, that is, the value of the knapsack is greater than or equal to the profit constraint.

In their study they implemented the knapsack decision problem in the form of the task presented in Fig 5c. In their task all instances had 6 items (*N* = 6) and *w*_*i*_, *v*_*i*_, *c* and *p* were integers. Each participant completed 72 trials (3 blocks of 24 trials with a rest period of 60s between blocks). Each trial presented a different instance of the KP. Trials had a time limit of 25 seconds and were *not* self-paced. A green circle at the center of the screen indicated the time remaining in each stage of the trial. During the first 3 seconds participants were presented with a set of items of different values and weights. Then, both capacity constraint and target profit were shown at the center of the screen for the remainder of the trial (22 seconds). No interactivity was incorporated into the task; that is, participants could not click on items. When the time limit was reached, participants were presented with the response screen where they could select one of two buttons: YES or NO. The time limit to respond was 2 seconds, and the inter-trial interval was 5 seconds. The order of instances and the sides of the YES/NO buttons on the response screen were randomized for each participant.

##### Instance sampling

It has been proposed that there exists a set of parameters 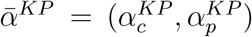 that captures the constrainedness of the problem, specifically 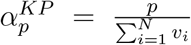 and 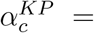 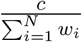 (*18*). These parameters characterize where typical instances are generally satisfiable (under-constrained region), where they are unsatisfiable (over-constrained region) and where the probability of satisfiability is close to 50% (satisfiability threshold). Instance near the satisfiability threshold have been shown to be, on average, harder to solve (*18*).

Instances in Franco et al. (*22*) were selected such that 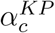 was fixed 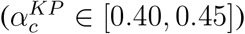 and the instance constrainedness varied according to 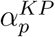. 18 instances were selected from the under-constrained region (*α*_*p*_ ∈ [0.35, 0.4]; *low TCC*) and 18 from the over-constrained region (*α*_*p*_ ∈ [0.85, 0.9]; *low TCC*). Additionally, 18 satisfiable instances and 18 unsatisfiable instances were sampled near the satisfiability threshold (*α*_*p*_ ∈ [0.6, 0.65]; *high TCC*).

Like for 3SAT and TSP, high TCC instances were selected such that they varied according to a specific measure of algorithmic complexity (number of propagations; see description of 3SAT sampling for details).

#### Procedure

After reading the plain language statement and providing informed consent, participants were instructed in the task and completed a practice session. Each experimental session lasted around 110 minutes. The tasks were programmed in Unity3D (*39*) and administered on a laptop.

Participants received a show-up fee of AUD 10 and additional monetary compensation based on accuracy. In the 3SAT, they additionally received AUD 0.6 for each correct instance submitted plus a bonus of AUD 0.31 per instance if all instances in the task were solved correctly. In the TSP, participants received 0.3 per correct instance submitted plus 0.14 per instance if all instances were solved correctly. In the KP task (*22*), participants received a show-up fee of AUD 10 and earned AUD 0.7 for each correct answer.

Note that the 3SAT and TSP tasks were self-paced (with time limits per trial), whereas the KP was not.

### Derivation of metrics

We estimated a collection of metrics based on the features of each instance and its solution space. We estimated one feature-space metric and several solution-space metrics. We first defined Typical-case complexity (TCC) according to the problem-parameter *α* for each task. Estimation of this metric is tightly related to the instance sampling procedure and its derivation is described in the previous section. Instances were sampled such that there was an equal number of instances with low and high TCC in each of the problems.

Once instances for the tasks were sampled, we estimated their solution-space metrics. Firstly, we calculated the *number of solution witnesses*, which corresponds to the number of state-space combinations (either paths, selection of items or switch-setups) that satisfy the constraints. For 3SAT, the number of solution witness was found using exhaustive search. For TSP instances, we used the Gecode algorithm (*40*) and allowed the algorithm to stop after finding 30,000 solution witnesses. This was done to reduce the computational requirements of solving an instance. 15 TSP instances reached the 30,000 maximum imposed. Given the variability in the number of witnesses in the TSP, the results on number witnesses are reported in logarithmic scale (natural logarithm).

Secondly, we estimated the instance complexity metric (IC) as the absolute value of the normalized difference between target value of the decision variant and the optimal value attainable of the corresponding optimization variant. In the KP, the optimization variant’s problem is to find the maximum value attainable given the weights, values and capacity. In the TSP, the optimization variant is to minimize the path traveled given a distance matrix. In the 3SAT, the optimization variant (MAXSAT) is to find the maximum number of satisfiable clauses given the Boolean formula presented. Explicitly, IC metrics are defined as follows:

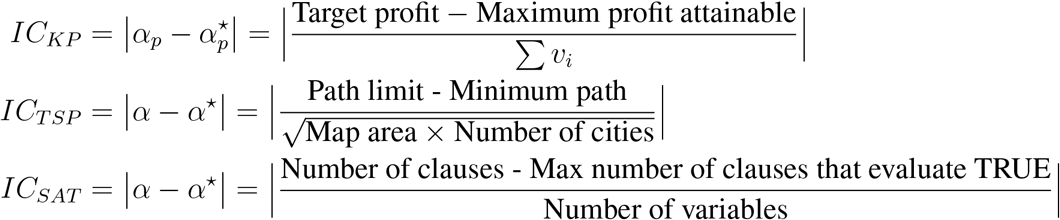

In order to estimate instance complexity (IC), the optimization variant of each instance needs to be solved. These optima were estimated using Gecode (*40*) for TSP and using the *RC2* algorithm from the ‘pysat’ python library (*41*) for 3SAT. For the KP we used the metrics estimated in Franco et al. (*22*).

### Statistical analysis

Python (version 3.6) was used to sample and solve instances. The R (version 3) programming language was used to analyze the behavioral data. All of the linear mixed models (LMM), generalized logistic mixed models (GLMM) and censored linear mixed models (CLMM) included random effects on the intercept for participants (unless otherwise stated). Different models were selected according to the data structure. GLMM were used for models with binary dependent variables, LMM were used for continuous dependent variables and CLMM were used for censored continuous dependent variables (e.g., time-on-task).

All the models were fitted using a Bayesian framework implemented using the probabilistic programming language Stan via the R package ‘brms’ (*42*). Default priors were used. All population-level effects of interest had uninformative priors; i.e., an improper flat prior over the reals. Intercepts had a student-t prior with 3 degrees of freedom and a scale parameter that depended on the standard deviation of the response after applying the link function. The student-t distribution was centered around the mean of the dependent variable. Sigma values, in the case of Gaussian-link models, had a half student-t prior (restricted to positive values) with 3 degrees of freedom and a scale parameter that depended on the standard deviation of the response after applying the link function. Standard deviation of the participant-level intercept parameters had a half student-t prior that was scaled in the same way as the sigma prior.

Each of the models presented was estimated using four Markov chains. The number of iterations per chain was by default set to 2000. This parameter was adjusted to 4000 on some models to ensure convergence. Convergence was verified using the convergence diagnostic 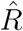. All models presented reach an 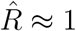.

Statistical tests were performed based on the 95% credible interval estimated using the highest density interval (HDI) of the posterior distributions calculated via the R package ‘parameters’ (*43*). For each statistical test we report both the median (*β*_0.5_) of the posterior distribution and its corresponding credible interval (*HDI*_0.95_).

For the knapsack task, we report the statistical results from (*22*) if available and are, here, reported as effect estimates (*β*) and P-Values (*P*). Otherwise we used the data available at the OSF (project: https://doi.org/10.17605/OSF.IO/T2JV7) to run statistical tests on the behavioral data. These tests were performed and reported following the same Bayesian approach used for the TSP and 3SAT analysis.

Some trials and participants were excluded due to different reasons. In the 3SAT task, two participants were omitted from the analysis given that their accuracy (close to 50%) differed significantly from the group. Additionally, 10 trials (from 9 participants) were omitted given that no answer was given. One participant was excluded from the time-on-task analysis since they never advanced to the response screen before the time limit. In the TSP, one participant was excluded from the analysis given that they did not understand the instructions. This was determined during the course of the experiment. Additionally 9 trials (from 8 participants) were omitted given that no answer was selected. Finally, in the knapsack task, 13 trials (from 8 participants) were excluded in which no response was made.

## Acknowledgments

The authors thank Elizabeth Bowman for her support of the laboratory experiments.

## Data and materials availability

The behavioral data and the data analysis code are both available at the Open Science Framework. The 3SAT and TSP tasks are also available there (project: https://osf.io/tekqa/).

## Author contributions

CM, JPF, KD, PB and NY designed the study; NY and JPF performed the instance selection; KD and JPF programmed the experimental tasks; KD ran a pilot version of this study; JPF performed data collection and analysis; JPF, CM, KD, NY and PB wrote the manuscript.

## Supplementary Materials

**S1 Appendix. Satisfiability and TCC.** Joint effect of satisfiability and TCC on human performance.

**S2 Appendix. Number of clicks.** The effect of instance properties on search length.

**S3 Appendix. Summary statistics.**

**S4 Appendix. TCC and the number of witnesses.** The number of witnesses drive the effect of TCC on accuracy in satisfiable instances.

**S5 Appendix. Explanatory power of IC in 3SAT.** Confounding factors.

**Figure S1 Number of Clicks.**

**Table S1 Human accuracy in the Boolean satisfiability task.**

**Table S2 Human accuracy in the traveling salesperson task.**

**Table S3 Time-on-task in the Boolean satisfiability task.**

**Table S4 Time-on-task in the traveling salesperson task.**

**Table S5 Human accuracy and the number of solution witnesses.**

**Table S6 Human accuracy in the knapsack task.**

**Table S7 Number of clicks in the Boolean satisfiability task.**

**Table S8 Number of clicks in the traveling salesperson task.**

**Table S9 Human performance and the number of clauses in SAT.**

