## Supplementary material for "Task-independent metrics of computational hardness predict human cognitive performance"

### Supplementary Materials for:
#### Task-independent metrics of computational hardness predict human cognitive performance

Juan P. Franco, Karlo Doroc, Nitin Yadav,
Peter Bossaerts, Carsten Murawski*

#### This PDF file includes:

**S1 Appendix. Satisfiability and TCC.** Joint effect of satisfiability and TCC on human performance.

**S2 Appendix. Number of clicks.** The effect of instance properties on search length.

**S3 Appendix. Summary statistics.**

**S4 Appendix. TCC and the number of witnesses.** The number of witnesses drive the effect of TCC on accuracy in satisfiable instances.

**S5 Appendix. Explanatory power of IC in 3SAT.** Confounding factors.

**Figure S1 Number of Clicks.**

**Table S1 Human accuracy in the Boolean satisfiability task.**

**Table S2 Human accuracy in the traveling salesperson task.**

**Table S3 Time-on-task in the Boolean satisfiability task.**

**Table S4 Time-on-task in the traveling salesperson task.**

**Table S5 Human accuracy and the number of solution witnesses.**

**Table S6 Human accuracy in the knapsack task.**

**Table S7 Number of clicks in the Boolean satisfiability task.**

**Table S8 Number of clicks in the traveling salesperson task.**

**Table S9 Human performance and the number of clauses in SAT.**

### S1 Appendix: Satisfiability and TCC

In the results section we showed that TCC has a detrimental effect on accuracy and time-on-task. Moreover, we see that more time is spent solving unsatisfiable instances. Interestingly, our results suggest that the effect of satisfiability on accuracy is modulated by the problem at hand. In order to study more closely how these generic features make instances hard for humans to solve across different problems, we explored whether TCC and satisfiability interact with each other to make instances harder.

We first explore the interaction effect between TCC and satisfiability on accuracy. Accuracy was not affected by satisfiability in low TCC instances in all three problems (TSP: $\beta_{0.5} = -0.16$, $HDI_{0.95} = [-0.85, 0.55]$, Table S2 Model 4; 3SAT: $\beta_{0.5} = -0.52$, $HDI_{0.95} = [-1.27, 0.11]$, Table S1 Model 4; KP: $\beta = -0.250$, $P = 0.355$, (*1*); marginal effect of satisfiability, GLMM). Moreover, in line with the previous KP study, we found a negative effect of TCC in both satisfiable and unsatisfiable instances for both problems considered (TSP: $\beta^{sat}_{0.5} = -2.07$, $HDI^{sat}_{0.95} = [-2.64, -1.55]$, $\beta^{unsat}_{0.5} = -2.16$, $HDI^{unsat}_{0.95} = [-2.74, -1.62]$, Table S2 Model 4; 3SAT: $\beta^{sat}_{0.5} = -2.06$, $HDI^{sat}_{0.95} = [-2.56, -1.59]$, $\beta^{unsat}_{0.5} = -0.77$, $HDI^{unsat}_{0.95} = [-1.49, -0.13]$, Table S1 Model 4; the effect of TCC on accuracy for satisfiable and unsatisfiable instances, respectively, GLMM). Interestingly, in the 3SAT problem we found that the reduction in accuracy due to TCC was larger for satisfiable instances ($\beta_{0.5} = -1.29$, $HDI_{0.95} = [-2.11, -0.46]$, interaction effect of TCC and satisfiability on accuracy, GLMM; Table S1 Model 4). In contrast, in the TSP, as with the KP, the size of the effect of TCC on accuracy was similar for both satisfiable and unsatisfiable instances ($\beta_{0.5} = 0.10$, $HDI_{0.95} = [-0.65, 0.90]$, interaction effect of TCC and satisfiability on accuracy, GLMM; Table S2 Model 4). This suggests that, unlike in the KP and TSP, in the 3SAT there is an interaction effect between TCC and satisfiability on accuracy, which makes satisfiable instances with high TCC harder than the rest.

When analyzing how satisfiability affected time-on-task for different levels of TCC we found different results across problems. In the TSP there was no interaction effect on time between satisfiability and TCC ($\beta_{0.5} = 0.002$, $HDI_{0.95} = [-0.054, 0.058]$, interaction effect of satisfiability and TCC on time-on-task, CLMM; Table S4 Model 7), meaning that both properties had independent effects on time-on-task. In contrast, in 3SAT the effect of TCC was modulated by satisfiability in such a way that there was no effect of TCC when the instance was unsatisfiable ($\beta_{0.5} = 0.022$, $HDI_{0.95} = [-0.016, 0.063]$, marginal effect of TCC on time-on-task for unsatisfiable instances, CLMM; Table S3 Model 8). In summary, we only found an interaction effect between satisfiability and TCC in the 3SAT. This was the case for both accuracy and time-on-task. These results suggest that satisfiability might have a differential effect on performance across different problems, including the way it interacts with TCC to make instances hard.

### S2 Appendix: Number of clicks

In order to validate whether the effect of the complexity metrics on time-on-task could be extended to other aspects of cognitive effort, we explored their effect on the number of clicks participants performed in each trial. The number of clicks is related to the extent in which the problem state-space is explored. In the 3SAT, the state-space consists of all possible on-off switch setups ($2^5$ possible combinations) while in the TSP the state-space consists of all possible ordered path selections ($2^{\binom{20}{2}} = 2^{190}$ possible combinations). Arguably, participants search the state-space by clicking on different state combinations in order to decide whether an instance is satisfiable or not. Differences in the quantity of clicks used to solve an instance can shed light into the cognitive effort exerted in exploring the state-space (under the assumption that the state-space is explored by clicking on elements in the task).

This analysis was performed for TSP and 3SAT. In both tasks, participants had the opportunity to click on cities or literals throughout the trial, whereas in the KP clicking on items was not possible. Note that while for the TSP there was no limit in the number of clicks, in the 3SAT participants were only allowed to make a maximum of 20 clicks per trial. The purpose of this limit was to discourage participants from using a trial-and-error strategy to solve the instances. Overall, the limit was reached in $11.5\%$ of the trials.

We investigated whether the generic complexity metrics could capture differences in the number of clicks. We found that participants performed more clicks on instances with high TCC, compared to low TCC, in 3SAT and TSP (TSP: $\beta_{0.5} = 1.66$, $HDI_{0.95} = [1.01, 2.33]$, GLMM Table S8 Model 1; 3SAT: $\beta_{0.5} = 1.88$, $HDI_{0.95} = [1.23, 2.54]$, CLMM, Table S7 Model 1; effect of TCC on number of clicks; Fig S1c). Additionally, less clicks were performed on satisfiable instances compared to unsatisfiable ones (TSP: $\beta_{0.5} = -2.48$, $HDI_{0.95} = [-3.13, -1.85]$, LMM Table S8 Model 2; 3SAT: $\beta_{0.5} = -7.41$, $HDI_{0.95} = [-7.95, -6.88]$, CLMM, Table S7 Model 2; effect of satisfiability on number of clicks; Fig S1b). When exploring how these two metrics jointly affected the length of search, we found that both effects were still significant when controlling for each other in the TSP (Table S8 Model 3; Fig S1b). However, in 3SAT the positive effect of TCC on the number of clicks was only present in satisfiable instances (Table S7 Model 3).

We then studied how the solution-space complexity metrics affected the length of search. Among satisfiable instances a higher number of witnesses was related to a lower amount of clicks (TSP: $\beta_{0.5} = -0.41$, $HDI_{0.95} = [-0.54, -0.26]$, LMM Table S8 Model 5; 3SAT: $\beta_{0.5} = -0.59$, $HDI_{0.95} = [-0.69, -0.49]$, CLMM, Table S7 Model 5; effect of number of witnesses on the number of clicks; Fig S1c). Additionally, we found that a higher IC value was related to lower number of clicks in the TSP and in unsatisfiable 3SAT instances (TSP: $\beta_{0.5} = -10.22$, $HDI_{0.95} = [-13.87, -6.37]$, LMM, Table S8 Model 4;

3SAT: $\beta_{0.5} = -29.88$, $HDI_{0.95} = [-54.52, -6.63]$, CLMM, Table S7 Model 4); effect of IC on number of clicks; Fig S1b). We excluded satisfiable 3SAT instances from the analysis since we are unable to disentangle the effect of IC and satisfiability; all 3SAT satisfiable instances have an $IC = 0$. In the TSP we investigated the joint effect of IC and satisfiability on the number of clicks and found that the effects were still significant when controlling for each other and that there was no interaction effect between the variables ($\beta_{0.5} = -5.95$, $HDI_{0.95} = [-13.43, 1.78]$, interaction effect between IC and satisfiability, LMM, Table S8 Model 6).

Overall, these finding closely follow those found for time-on-task. Indeed, the only significant difference is that the negative effect of IC on number of clicks for unsatisfiable 3SAT instances is statistically significant. This suggests that the related dimension of cognitive effort, namely the extent to which the state-space is searched, can be elucidated by the intrinsic properties of an instance.

### S3 Appendix: Summary statistics

In this section we present summary statistics of the behavioral data for each of the tasks. These statistics exclude some observations as described in the Statistical Analysis section in Materials and Methods.

#### Boolean satisfiability task

On average, participants chose the 'YES' option in 45% of trials ($\min = 28\%$, $\max = 61\%$). Accuracy did not vary during the course of the task ($\beta_{0.5} = 0.001$, $HDI_{0.95} = [-0.008, 0.011]$, main effect of trial number on accuracy, generalized logistic mixed model

(GLMM); Table S1 Model 1), suggesting that neither experience with the task nor mental fatigue affected task accuracy. However, time spent did vary throughout the task. As the task progressed they spent on average less time on a trial ($\beta_{0.5} = -0.005$, $HDI_{0.95} = [-0.006, -0.004]$, main effect of trial number on time-on-task —as a proportion of the maximum possible time—, censored linear mixed effects model (CLMM); Table S3 Model 1). Overall, participants reached the maximum time allotted (110 seconds) 16% of trials.

#### Traveling salesperson task

On average, participants chose the 'YES' option in 50% of trials ($\mathrm{min} = 35\%$, $\mathrm{max} = 60\%$). Consistent with our results for the 3SAT, accuracy did not vary during the course of the task ($\beta_{0.5} = 0.00$, $HDI_{0.95} = [-0.003, 0.013]$, main effect of trial number on accuracy, GLMM; Table S2 Model 1), and participants spent less time on a trial as they progressed ($\beta_{0.5} = -0.002$, $HDI_{0.95} = [-0.003, -0.001]$, main effect of trial number on time-on-task —as a proportion of maximum possible time—), CLMM; Table S4 Model 1). Overall, participants reached the limit of 40 seconds 42.9% of trials.

#### Knapsack decision task

On average, participants chose the 'YES' option in 48.1% of trials ($\mathrm{min} = 0.32$, $\mathrm{max} = 0.60$, $SD = 0.06$). Accuracy did not vary during the course of the task ($\beta = 0.005$, $P = 0.196$, main effect of trial number on accuracy, GLMM; (*1*)).

### S4 Appendix: TCC and the number of witnesses

It is feasible that the effect of TCC on accuracy, in satisfiable instances, is driven by the number of witnesses. After all, TCC is constructed as a mapping from constrainedness ($\alpha$) to computational hardness, and the number of solution witnesses captures directly the 'constrainedness' of the solutions space of a single instance. We thus examined the link between these two metrics. As expected, we found that the number of witnesses of low TCC instances was significantly higher than that of instances with high TCC in all three problems ($P_{SAT} < 0.001$, $P_{TSP} < 0.001$, $P_{KP} < 0.001$, p-values of unpaired t-tests).

To test whether the effect on accuracy of TCC (in satisfiable instances) is driven by the number of witnesses we studied the effect of TCC on accuracy while controlling for the number of witnesses. In line with our conjecture, we found that once we controlled for the number of witnesses the marginal effect of TCC on accuracy was not significant on all three problems (3SAT: $\beta_{0.5} = 0.47$, $HDI_{0.95} = [-0.30, 1.22]$; TSP: $\beta_{0.5} = -0.12$, $HDI_{0.95} = [-0.84, 0.68]$; KP: $\beta_{0.5} = 0.17$, $HDI_{0.95} = [-0.57, 0.90]$; marginal effect of TCC on accuracy, GLMM; Table S5 Models 2,5,8). We studied further this relation and tested whether there was an interaction effect of TCC and the number of witnesses on accuracy. The results were different across problems. We found a significant interaction in the KP, an inconclusive result in the 3SAT and a non-significant results in the TSP (KS: $\beta_{0.5} = 0.54$, $HDI_{0.95} = [0.26, 0.81]$; 3SAT: $\beta_{0.5} = 0.56$, $HDI_{0.95} = [-0.00, 1.22]$; TSP: $\beta_{0.5} = -0.26$, $HDI_{0.95} = [-0.66, 0.15]$; interaction effect between TCC and number of witnesses on accuracy, GLMM; Table S5 Models 3,6,9). Taken together, these results suggest that the effect of TCC on accuracy is, at least partially, driven by the number of witnesses. However, on some problems, TCC might affect human accuracy through other mechanisms as well.

### S5 Appendix: Explanatory power of IC in 3SAT

IC explains less variability in accuracy in 3SAT than in the other two tasks. One possible explanation is that, unlike in TSP and KP, in 3SAT the IC metric takes a values of zero ($IC = 0$) when the instance is satisfiable; by definition an instance is only satisfiable if the maximum number of clauses set to TRUE is equal to the number of clauses in the instance (i.e., $IC = 0$). This entails that IC would only be able to explain differences in accuracy in unsatisfiable instances, which (given our sampling procedure) amount to half of the instances used in the task.

We tested whether this characteristic was driving the lower explanatory power of IC on accuracy in our sample. To do this, we first restricted our analysis to unsatisfiable instances, where the explanatory power should be unaffected by this characteristic of the instances sampled. We found that the explanatory power for these instances was still low ($R^2 = 0.001$ for unsatisfiable 3SAT instances). Moreover, for this set of instances the positive relation between IC and accuracy was not significant in 3SAT, in contrast to the KP and TSP, where the positive relation between IC and accuracy was significant in unsatisfiable instances (KP: $\beta_{0.5} = 13.48$, $HDI_{0.95} = [10.11, 17.30]$, Table S6 Model 3; TSP: $\beta_{0.5} = 20.10$, $HDI_{0.95} = [15.39, 24.96]$, Table S2 Model 8; 3SAT: $\beta_{0.5} = 9.81$, $HDI_{0.95} = [-14.59, 33.94]$, Table S1 Model 7; the effect of IC on accuracy for unsatisfiable instances, GLMM).

Overall, these results indicate that IC is able to explain variance in accuracy across instances, but to a lesser degree in 3SAT. This is even, after considering the characteristics of the sampled 3SAT instances in which this is tested. Moreover, the fact that more proportion of the variance is explained by IC when considering all 3SAT instances ($R^2 = 0.16$) compared to only unsatisfiable instances ($R^2 = 0.001$) suggests that the effect of IC on accuracy in the 3SAT might be incongruously driven by satisfiability. However, we are unable to properly disentangle the effect of IC and satisfiability given that all 3SAT satisfiable instance have $IC = 0$.

### S6 Appendix: Supplementary Figures and Tables

Figure 1: **Number of Clicks. (a) Satisfiability and TCC.** Median number of clicks performed while solving an instance before submitting an answer. Each colored dot represents an instance of a problem. **(b) IC.** Each green and orange shape represent the median number of clicks for each instance of TSP and 3SAT problems. The blue lines represents the marginal effect of IC (LMM, Table S8 Model 4, CLMM Table S7 Model 4). Satisfiable 3SAT instances are excluded from the 3SAT model since we are unable to disentangle the effect of IC and satisfiability. Each black dot corresponds to the number of clicks by a single participant on a particular instance. The range of number of clicks presented for the TSP ($[10, 50]$) contains more than $98\%$ of observations. **(c) Number of solution witnesses.** Each green shape represents the median time-on-task per instance. The blue line represents the marginal effect of the number of solution witnesses (LMM, Table S8 Model 5 and CLMM, Table S7 Model 5). *The box-plots represent the median, the interquartile range (IQR) and the whiskers extend to a maximum length of 1.5*IQR.*

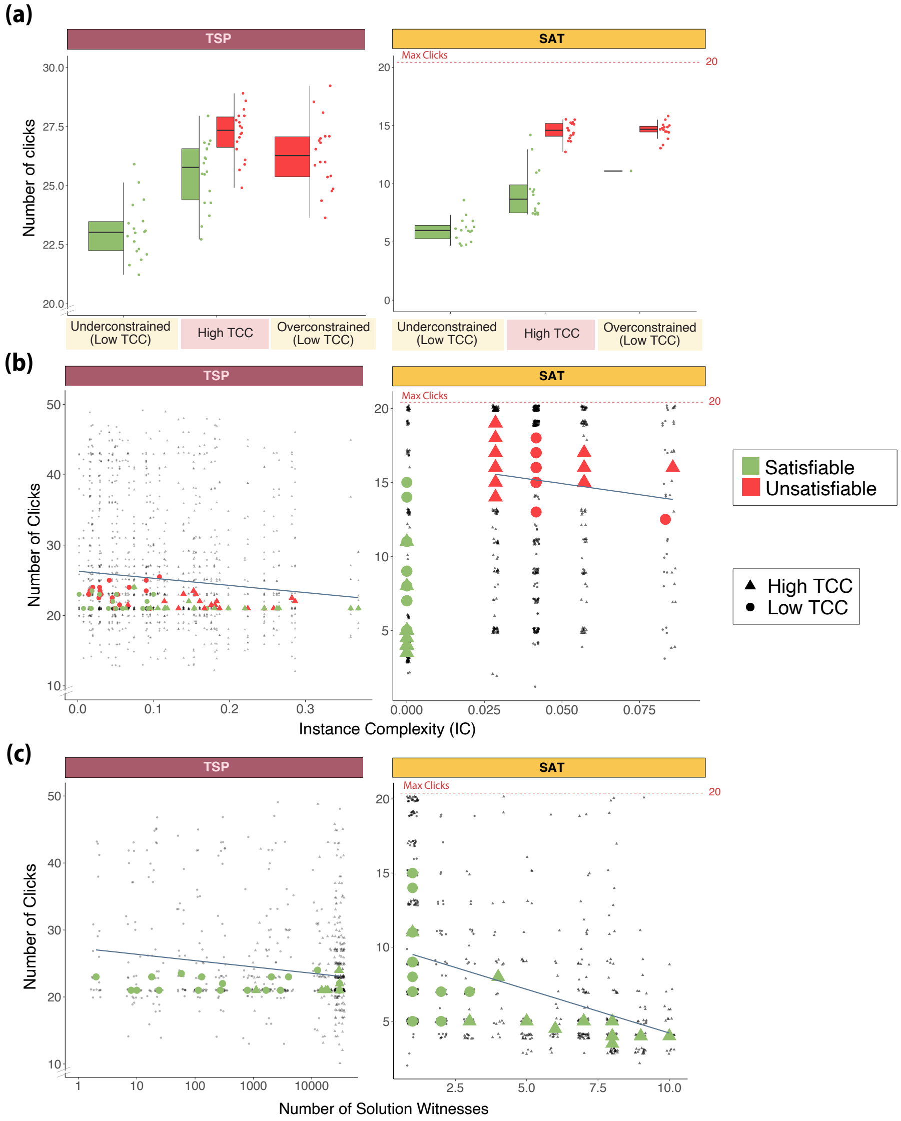

Table 1: **Human accuracy in the Boolean satisfiability task.** Logistic regressions with random intercept effects for participants relating the accuracy on an instance and trial number (1), typical-case complexity (TCC) (2), constrainedness region (3), TCC and satisfiability (4), time-on-task (5), instance complexity (IC) (6), IC on unsatisfiable instances (7) as well as satisfiability (8). *Parameter estimates correspond to the median of the posterior distribution ($\beta_{0.5}$) and the 95% HDI credible interval ($HDI_{0.95}$). ELPD denotes the expected log posterior predictive density.*

| | Dependent variable: Human accuracy | | | | | | | |
|---|---|---|---|---|---|---|---|---|
| | (1) | (2) | (3) | (4) | (5) | (6) | (7) | (8) |
| Trial number | 0<br>[-0.01,0.01] | | | | | | | |
| TCC | | -1.58<br>[-1.95,-1.2] | | -0.77<br>[-1.49,-0.13] | | | | |
| Overconstrained | | | 1.39<br>[0.93,1.86] | | | | | |
| Underconstrained | | | 1.82<br>[1.31,2.4] | | | | | |
| Satisfiability | | | | -0.52<br>[-1.27,0.11] | | | | -1.35<br>[-1.73,-0.99] |
| TCC:Satisfiability | | | | -1.29<br>[-2.11,-0.46] | | | | |
| Time-on-task | | | | | -0.02<br>[-0.02,-0.01] | | | |
| IC | | | | | | 30.3<br>[21.95,39.24] | 9.81<br>[-14.59,33.94] | |
| Intercept | 1.95<br>[1.59,2.26] | 2.99<br>[2.59,3.43] | 1.41<br>[1.13,1.75] | 3.33<br>[2.73,3.97] | 3.15<br>[2.53,3.73] | 1.51<br>[1.23,1.8] | 2.99<br>[1.58,4.29] | 2.84<br>[2.43,3.22] |
| Observations | 1398 | 1398 | 1398 | 1398 | 1335 | 1398 | 675 | 1398 |
| ELPD | -533.3 | -493.27 | -493.42 | -456.8 | -487.73 | -505.05 | -138.46 | -503.36 |

Table 2: **Human accuracy in the traveling salesperson task.** Logistic regressions with random intercept effects for participants relating the accuracy on an instance and trial number (1), typical-case complexity (TCC) (2), constrainedness region (3), TCC and satisfiability (4), time-on-task (5), satisfiability (6), instance complexity (IC) (7), as well as IC on unsatisfiable instances (8). *Parameter estimates correspond to the median of the posterior distribution ($\beta_{0.5}$) and the 95% HDI credible interval ($HDI_{0.95}$). ELPD denotes the expected log posterior predictive density.*

| | Dependent variable: Human accuracy | | | | | | | |
|---|---|---|---|---|---|---|---|---|
| | (1) | (2) | (3) | (4) | (5) | (6) | (7) | (8) |
| Trial number | 0<br>[0,0.01] | | | | | | | |
| TCC | | -2.1<br>[-2.5,-1.73] | | -2.16<br>[-2.74,-1.62] | | | | |
| Overconstrained | | | 2.18<br>[1.65,2.73] | | | | | |
| Underconstrained | | | 2.05<br>[1.56,2.63] | | | | | |
| Satisfiability | | | | -0.16<br>[-0.85,0.55] | | -0.06<br>[-0.34,0.22] | | |
| TCC:Satisfiability | | | | 0.1<br>[-0.65,0.9] | | | | |
| Time-on-task | | | | | -0.1<br>[-0.13,-0.08] | | | |
| IC | | | | | | | 21.13<br>[17.63,24.91] | 20.1<br>[15.39,24.96] |
| Intercept | 1.66<br>[1.43,1.94] | 3.19<br>[2.83,3.58] | 1.09<br>[0.91,1.3] | 3.29<br>[2.8,3.84] | 5.36<br>[4.39,6.34] | 1.81<br>[1.59,2.03] | 0.14<br>[-0.14,0.4] | 0.52<br>[-0.21,1.29] |
| Observations | 1575 | 1575 | 1575 | 1575 | 1575 | 1575 | 1575 | 787 |
| ELPD | -656.28 | -578.48 | -579.52 | -580.43 | -612.95 | -656.93 | -534.35 | -241.48 |

Table 3: **Time-on-task in the Boolean satisfiability task.** Censored linear regressions with random intercept effects for participants relating the time spent on an instance (as a proportion of the maximum time allotted on each trial) and trial number (1), typical-case complexity (TCC) (2), constrainedness region (3), satisfiability (4), number of solution witnesses (5), instance complexity (IC) (6), IC on unsatisfiable instances (7), as well as TCC and satisfiability (8). *Parameter estimates correspond to the median of the posterior distribution ($\beta_{0.5}$) and the 95% HDI credible interval ($HDI_{0.95}$). ELPD denotes the expected log posterior predictive density.*

| | Dependent variable: Time-on-task | | | | | | | |
|---|---|---|---|---|---|---|---|---|
| | (1) | (2) | (3) | (4) | (5) | (6) | (7) | (8) |
| Trial number | -0.01<br>[-0.01,0] | | | | | | | |
| TCC | | 0.15<br>[0.12,0.18] | | | | | | 0.02<br>[-0.02,0.06] |
| Overconstrained | | | 0.07<br>[0.04,0.11] | | | | | |
| Underconstrained | | | -0.35<br>[-0.39,-0.32] | | | | | |
| Satisfiability | | | | -0.32<br>[-0.35,-0.29] | | | | -0.42<br>[-0.46,-0.38] |
| No. of witnesses | | | | | -0.04<br>[-0.05,-0.04] | | | |
| IC | | | | | | 6.04<br>[5.41,6.7] | -0.56<br>[-1.91,0.78] | |
| TCC:Satisfiability | | | | | | | | 0.21<br>[0.16,0.27] |
| Intercept | 0.68<br>[0.65,0.72] | 0.51<br>[0.4,0.62] | 0.65<br>[0.54,0.75] | 0.74<br>[0.64,0.84] | 0.59<br>[0.51,0.69] | 0.45<br>[0.35,0.57] | 0.77<br>[0.63,0.91] | 0.73<br>[0.62,0.83] |
| Observations | 1335 | 1335 | 1335 | 1335 | 691 | 1335 | 644 | 1335 |
| ELPD | -683.08 | -456.79 | -268.31 | -296.9 | -15.46 | -340.59 | -128.73 | -224.94 |

Table 4: **Time-on-task in the traveling salesperson task.** Censored linear regressions with random intercept effects for participants relating the time spent on an instance (as a proportion of the maximum time allotted on each trial) and trial number (1), typical-case complexity (TCC) (2), constrainedness region (3), satisfiability (4), number of solution witnesses (scaled via natural logarithm) (5), instance complexity (IC) (6), as well as TCC and satisfiability (7). *Parameter estimates correspond to the median of the posterior distribution ($\beta_{0.5}$) and the 95% HDI credible interval ($HDI_{0.95}$). ELPD denotes the expected log posterior predictive density.*

| | Dependent variable: Time-on-task | | | | | | |
|---|---|---|---|---|---|---|---|
| | (1) | (2) | (3) | (4) | (5) | (6) | (7) |
| Trial number | 0.00<br>[0.00,0.00] | | | | | | |
| TCC | | 0.12<br>[0.09,0.15] | | | | | 0.12<br>[0.08,0.16] |
| Overconstrained | | | -0.02<br>[-0.06,0.01] | | | | |
| Underconstrained | | | -0.2<br>[-0.23,-0.16] | | | | |
| Satisfiability | | | | -0.17<br>[-0.2,-0.15] | | | -0.17<br>[-0.21,-0.14] |
| No. of witnesses (ln) | | | | | -0.02<br>[-0.03,-0.02] | | |
| IC | | | | | | -0.74<br>[-0.9,-0.58] | |
| TCC:Satisfiability | | | | | | | 0.00<br>[-0.05,0.06] |
| Intercept | 0.97<br>[0.87,1.08] | 0.85<br>[0.75,0.95] | 0.97<br>[0.87,1.06] | 1.00<br>[0.89,1.1] | 0.99<br>[0.89,1.09] | 1.00<br>[0.9,1.11] | 0.94<br>[0.84,1.04] |
| Observations | 1575 | 1575 | 1575 | 1575 | 788 | 1575 | 1575 |
| ELPD | -515.38 | -499.04 | -460.55 | -459.22 | -189.73 | -491.96 | -426.44 |

Table 5: **Human accuracy and the number of solution witnesses.** Logistic regressions with random intercept effects for participants with accuracy as dependent variable. The data included on each regression is comprised of the satisfiable instances of one of the three tasks considered: 3SAT (1-3), TSP (4-6) and KP (7-9). Regressions (1), (4) and (7) include the the number of witnesses alone as regressor (the number of witnesses for the TSP is scaled via natural logarithm). Models (2), (5) and (8) include TCC, additionally, as regressor. Models (3), (5) and (8) include the interaction between TCC and number of witnesses as well. *Parameter estimates correspond to the median of the posterior distribution ($\beta_{0.5}$) and the 95% HDI credible interval ($HDI_{0.95}$). ELPD denotes the expected log posterior predictive density.*

| | Dependent variable: Human accuracy | | | | | | | | |
|---|---|---|---|---|---|---|---|---|---|
| | 3SAT | | | TSP | | | KP | | |
| | (1) | (2) | (3) | (4) | (5) | (6) | (7) | (8) | (9) |
| No. of witnesses | 0.62<br>[0.49,0.79] | 0.7<br>[0.51,0.91] | 0.63<br>[0.44,0.85] | | | | 0.26<br>[0.19,0.34] | 0.29<br>[0.17,0.41] | 0.11<br>[-0.03,0.25] |
| TCC | | 0.47<br>[-0.3,1.22] | -0.42<br>[-1.71,0.7] | | -0.12<br>[-0.84,0.68] | 2.24<br>[-1.5,6.13] | | 0.17<br>[-0.57,0.9] | -1.77<br>[-2.95,-0.44 |
| TCC:No. of witnesses | | | 0.56<br>[0,1.22] | | | | | | 0.54<br>[0.26,0.81] |
| No. of witnesses (ln) | | | | 0.45<br>[0.37,0.53] | 0.44<br>[0.33,0.54] | 0.68<br>[0.25,1.03] | | | |
| TCC:No. of witnesses (ln) | | | | | | -0.26<br>[-0.66,0.15] | | | |
| Intercept | -0.02<br>[-0.62,0.53] | -0.52<br>[-1.58,0.47] | -0.21<br>[-1.19,0.93] | -1.07<br>[-1.78,-0.46] | -0.92<br>[-2.16,0.21] | -3.2<br>[-6.83,0.73] | 0.41<br>[-0.04,0.83] | 0.22<br>[-0.7,1.22] | 1.55<br>[0.35,2.78] |
| Observations | 723 | 723 | 723 | 788 | 788 | 788 | 716 | 716 | 716 |
| ELPD | -258.83 | -259.54 | -258.59 | -244.8 | -245.4 | -245.76 | -303.21 | -303.92 | -297.45 |

Table 6: **Human accuracy in the knapsack task.** Logistic regressions with random intercept effects for participants relating the accuracy on an instance and satisfiability (1), instance complexity (IC) (2), and IC on only unsatisfiable instances (3). *Parameter estimates correspond to the median of the posterior distribution (*$\beta_{0.5}$*) and the 95% HDI credible interval (*$HDI_{0.95}$*). ELPD denotes the expected log posterior predictive density.*

| | Dependent variable: Human accuracy | | |
|---|---|---|---|
| | (1) | (2) | (3) |
| Satisfiability | -0.29 | | |
| | [-0.57,0.01] | | |
| IC | | 9.05 | 13.48 |
| | | [7.2,11.02] | [10.11,17.3] |
| Intercept | 1.79 | 0.59 | 0.47 |
| | [1.51,2.1] | [0.28,0.92] | [0.08,0.92] |
| Observations | 1427 | 1427 | 711 |
| ELPD | -637.57 | -574.82 | -253.01 |

Table 7: **Number of clicks in the Boolean satisfiability task.** Censored linear regressions with random intercept effects for participants relating the number of clicks performed on an instance and typical-case complexity (TCC) (1), satisfiability (2), TCC and satisfiability (3), instance complexity (IC) on unsatisfiable instances (4), as well as the number of solution witnesses on satisfiable instances (5). *Parameter estimates correspond to the median of the posterior distribution ($\beta_{0.5}$) and the 95% HDI credible interval ($HDI_{0.95}$). ELPD denotes the expected log posterior predictive density.*

| | Dependent variable: Number of clicks | | | | |
|---|---|---|---|---|---|
| | (1) | (2) | (3) | (4) | (5) |
| TCC | 1.88<br>[1.23,2.54] | | 0.06<br>[-0.66,0.84] | | |
| Satisfiability | | -7.41<br>[-7.95,-6.88] | -8.8<br>[-9.51,-8.05] | | |
| TCC:Satisfiability | | | 2.91<br>[1.81,3.88] | | |
| IC | | | | -29.88<br>[-54.52,-6.63] | |
| No. of witnesses | | | | | -0.59<br>[-0.69,-0.49] |
| Intercept | 10.43<br>[9.19,11.67] | 15.15<br>[13.95,16.39] | 15.11<br>[13.76,16.4] | 16.4<br>[14.19,18.82] | 10.09<br>[9.23,10.96] |
| Observations | 1375 | 1375 | 1375 | 664 | 711 |
| ELPD | -4111.67 | -3825.19 | -3794.12 | -1645.16 | -2016.33 |

Table 8: **Number of clicks in the traveling salesperson task.** Linear regressions with random intercept effects for participants relating the number of clicks performed on an instance and typical-case complexity (TCC) (1), satisfiability (2), TCC and satisfiability (3), instance complexity (IC) (4), the number of solution witnesses (transformed via natural logarithm) on satisfiable instances (5), as well as IC and satisfiability (6). *Parameter estimates correspond to the median of the posterior distribution ($\beta_{0.5}$) and the 95% HDI credible interval ($HDI_{0.95}$). ELPD denotes the expected log posterior predictive density.*

| | Dependent variable: Number of clicks | | | | | |
|---|---|---|---|---|---|---|
| | (1) | (2) | (3) | (4) | (5) | (6) |
| TCC | 1.66<br>[1.01,2.33] | | 0.89<br>[0.05,1.83] | | | |
| Satisfiability | | -2.48<br>[-3.13,-1.85] | -3.23<br>[-4.12,-2.3] | | | -1.77<br>[-2.81,-0.65] |
| TCC:Satisfiability | | | 1.53<br>[0.2,2.77] | | | |
| IC | | | | -10.22<br>[-13.87,-6.37] | | -6.71<br>[-12.41,-0.87] |
| TCC:No. of witnesses (ln) | | | | | -0.41<br>[-0.54,-0.26] | |
| IC:Satisfiability | | | | | | -5.95<br>[-13.43,1.78] |
| Intercept | 24.21<br>[21.99,26.32] | 26.38<br>[24.09,28.56] | 26<br>[23.61,28.28] | 26.29<br>[24.04,28.61] | 27.31<br>[25.52,29.17] | 27.2<br>[24.87,29.46] |
| Observations | 1575 | 1575 | 1575 | 1575 | 788 | 1575 |
| ELPD | -5236.1 | -5220.82 | -5207.65 | -5234.03 | -2544.04 | -5207.35 |

Table 9: **Human performance and the number of clauses in SAT.** The effect of the number of clauses and TCC on human accuracy —logistic regression— (1), and time spent on an instance (as a proportion of the maximum time allotted on each trial) —linear regression— (2). Both regressions include random intercept effects for participants. *Parameter estimates correspond to the median of the posterior distribution ($\beta_{0.5}$) and the 95% HDI credible interval ($HDI_{0.95}$). ELPD denotes the expected log posterior predictive density.*

| Dependent variable: | Human accuracy | Time-on-task |
|---|---|---|
| | (1) | (2) |
| TCC | -1.59 | 0.12 |
| | [-1.94,-1.20] | [0.09,0.14] |
| Number of clauses | -0.02 | 0.02 |
| | [-0.05,0.01] | [0.02,0.02] |
| Intercept | 3.4 | 0.13 |
| | [2.65,4.25] | [0.03,0.25] |
| Observations | 1398 | 1335 |
| ELPD | -493.38 | -268.19 |
